# *C9orf72*-derived arginine-containing dipeptide repeats associate with axonal transport machinery and impede microtubule-based motility

**DOI:** 10.1101/835082

**Authors:** Laura Fumagalli, Florence L. Young, Steven Boeynaems, Mathias De Decker, Arpan R. Mehta, Ann Swijsen, Raheem Fazal, Wenting Guo, Matthieu Moisse, Jimmy Beckers, Lieselot Dedeene, Bhuvaneish T. Selvaraj, Tijs Vandoorne, Vanesa Madan, Marka van Blitterswijk, Denitza Raitcheva, Alexander McCampbell, Koen Poesen, Aaron D. Gitler, Phillip Koch, Pieter Vanden Berghe, Dietmar Rudolf Thal, Catherine Verfaillie, Siddharthan Chandran, Ludo Van Den Bosch, Simon L. Bullock, Philip Van Damme

**Author notes:** these authors contributed equally (Laura Fumagalli, Florence L. Young, Steven Boeynaems).

## Abstract

Hexanucleotide repeat expansions in the *C9orf72* gene are the most common genetic cause of amyotrophic lateral sclerosis (ALS) and frontotemporal dementia (FTD). How this mutation leads to these neurodegenerative diseases remains unclear. Here, we use human induced pluripotent stem cell-derived motor neurons to show that *C9orf72* repeat expansions impair microtubule-based transport of mitochondria, a process critical for maintenance of neuronal function. Cargo transport defects are recapitulated by treating healthy neurons with the arginine-rich dipeptide repeat proteins (DPRs) that are produced by the hexanucleotide repeat expansions. Single-molecule imaging shows that these DPRs perturb motility of purified kinesin-1 and cytoplasmic dynein-1 motors along microtubules *in vitro*. Additional *in vitro* and *in vivo* data indicate that the DPRs impair transport by interacting with both microtubules and the motor complexes. We also show that kinesin-1 is enriched in DPR inclusions in patient brains and that increasing the level of this motor strongly suppresses the toxic effects of arginine-rich DPR expression in a *Drosophila* model. Collectively, our study implicates an inhibitory interaction of arginine-rich DPRs with the axonal transport machinery in *C9orf72*-associated ALS/FTD and thereby points to novel potential therapeutic strategies.

## INTRODUCTION

A GGGGCC (G_4_C_2_) repeat expansion in the *C9orf72* gene is the most common genetic cause of both amyotrophic lateral sclerosis (ALS) and frontotemporal dementia (FTD) (*C9*-ALS/FTD) (1, 2). Since the association between the hexanucleotide repeat expansions (HREs) and these neurodegenerative diseases was discovered, three non-mutually exclusive pathological mechanisms have been proposed. The first is a loss-of-function scenario due to decreased expression of *C9orf72* transcript and protein observed in *C9*-ALS/FTD patients (1, 3). The second is an RNA gain-of-function mechanism caused by the accumulation of expanded repeat transcripts that sequester numerous RNA-binding proteins (4–9). The third proposed mechanism is a protein gain-of-function via the generation of pathological dipeptide repeat proteins (DPRs) originating from non-ATG mediated translation of the expanded repeat transcripts (10–13).

This repeat-associated non-ATG (RAN) translation occurs in all reading frames of sense and antisense transcripts resulting in five DPR proteins: poly-GR and poly-GA exclusively from the sense transcript, poly-PR and poly-PA exclusively from the antisense transcript, and poly-GP from both transcripts (10–13). DPRs are found in cytoplasmic inclusions in *C9*-ALS/FTD post-mortem brain and spinal cord tissue, and also have been detected in motor neurons differentiated from patient-derived induced pluripotent stem cells (iPSCs) (5,10,11,14–18). The arginine-rich DPRs – poly-PR and poly-GR – are potently toxic in numerous disease models (14,19–27), and have been shown to cause mitochondrial (28, 29) and endoplasmic reticulum stress (26), as well as disturbances in gene expression, RNA processing and translation (14,30–34), nucleocytoplasmic transport (21,22,25,35) and the dynamics of membrane-less organelles (36–38). Whether these effects fully explain the toxicity of arginine-rich DPRs remains unclear.

Of the several other genes that have been associated with ALS (39), three encode proteins important for microtubule-based cargo transport: the tubulin isotype α4a (40), the plus end-directed kinesin-1 motor KIF5A (41), and DCTN1, a component of the dynactin complex that activates the minus end-directed motor cytoplasmic dynein-1 (hereafter dynein) (42). The association of these mutations with ALS suggests that motor neurons, which are selectively targeted by the disease, are particularly reliant on efficient cargo trafficking due to their extended processes. This notion prompted us to investigate the involvement of microtubule-based transport in *C9*-ALS/FTD.

We observe that microtubule-based transport is impaired in iPSC-derived motor neurons from *C9*-ALS/FTD patients, and that treatment with arginine-rich DPRs elicits comparable defects in healthy neurons. Single-molecule imaging reveals that poly-GR and poly-PR impair the motility of purified kinesin-1 and dynein motors along microtubules *in vitro*. Analysis of physical interactions, including in iPSC-derived motor neurons and post-mortem patient brains, demonstrate association of arginine-rich DPRs with both microtubules and motor proteins. Finally, genetic analysis in *Drosophila* provides further evidence of a link between microtubule-based transport machinery and the toxic effects of arginine-rich DPRs. Collectively, our data strengthen the evidence that defective axonal cargo trafficking contributes to ALS pathogenesis and implicate inhibitory interactions of arginine-rich DPRs with the microtubule-based transport machinery in *C9*-ALS/FTD.

## RESULTS

### *C9orf72* hexanucleotide expansions impair microtubule-based transport in motor neurons

To investigate whether *C9*-ALS/FTD is associated with impaired microtubule-based transport we generated spinal motor neurons (sMNs) from fibroblast-derived iPSC lines from four *C9orf72* patients and three healthy controls (**Fig. S1, Fig. S2, Fig. S3** and **Table S1**). These *C9orf72* lines produced motor neurons with similar efficiency to control lines (**Fig. S3**). Whole-cell patch clamp recordings of evoked and spontaneous action potentials demonstrated functional maturation of sMNs with no differences in electrical activity between mutant *C9orf72* and control sMNs (**Fig. S4** and **Table S2**). Loss-of-function disease mechanisms have been proposed in *C9*-ALS/FTD (1, 3). To evaluate whether the repeat expansion results in reduction of *C9orf72* mRNA levels we performed digital droplet quantitative PCR. We found no changes in *C9orf72* transcript levels in the mutant cell lines compared to controls, although some subtle differences in the relative levels of the different transcript variants were observed (**Fig. S5A-D**). Using an enzyme-linked immunosorbent assay (ELISA), we could detect both poly-GA and poly-GP in all the *C9orf72* lines (**Fig. S5E**), confirming that RAN translation is occurring in these cells.

Defective microtubule-based transport of mitochondria in axons has been linked with a variety of neurodegenerative diseases (43), including ALS caused by other mutations (44–48). We therefore assessed motility of these organelles by live imaging of *C9orf72* and healthy control motor neuron lines incubated with the mitochondrial stain, Mitotracker. Kymograph-based analysis of neurons that had been differentiated for 38 days revealed that the proportion of mitochondria undergoing transport in neurites was reduced in the *C9orf72* lines (Fig. 1A, B and **Fig. S6A-D**). Transport was not significantly affected in younger neurons, revealing an age-related onset of the phenotype (Fig. 1C and **Fig. S6E-G**).

**Fig. 1:**
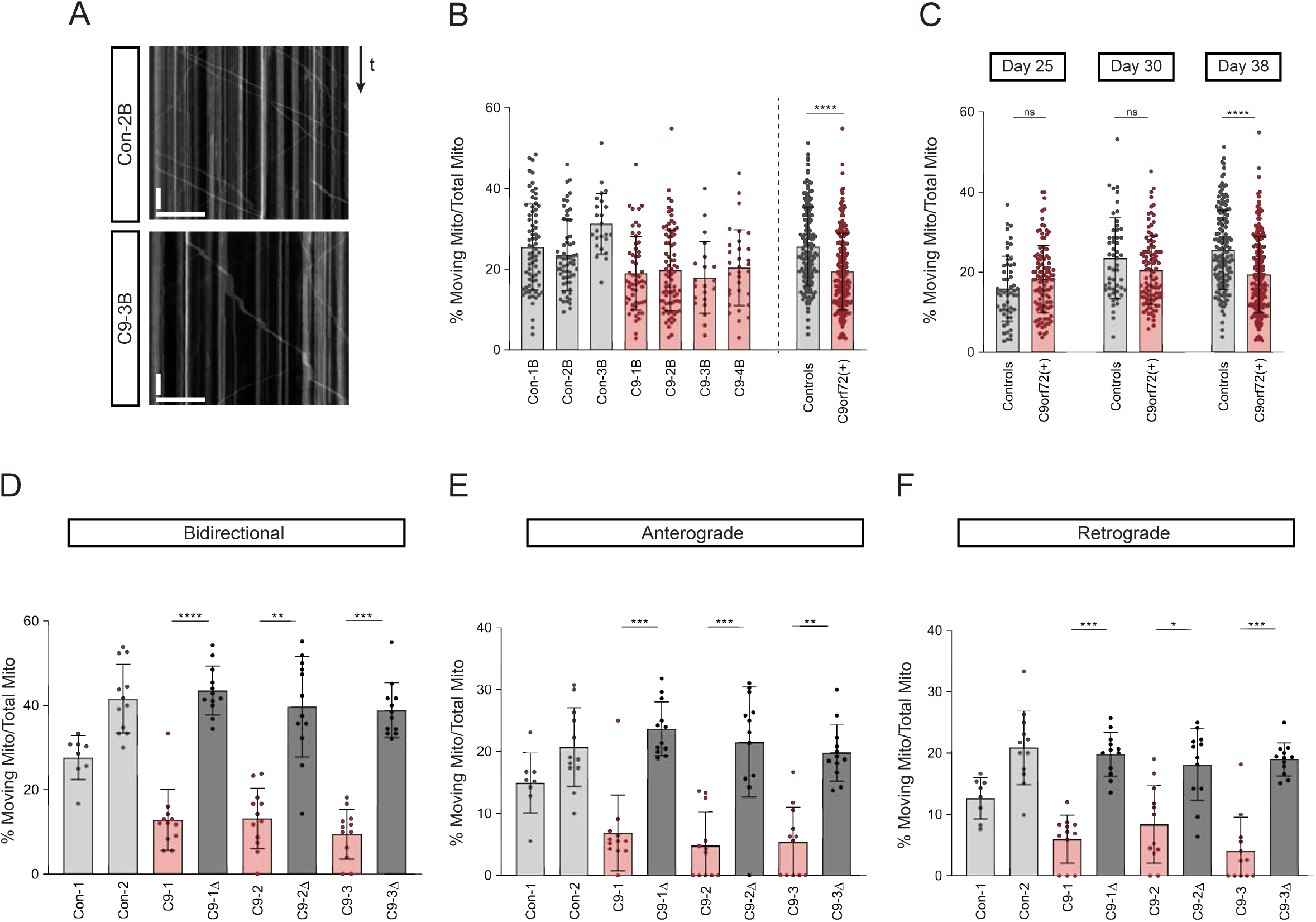
The *C9orf72* hexanucleotide repeat expansion causes deficits in mitochondria transport in sMNs. (A) Example kymographs (time-distance plots) from control and *C9orf72* sMNs showing MitoTracker-Red labeled stationary and moving mitochondria. Y-axis, time; x-axis, distance; scale bars, 50 s and 30 µm. Stationary and moving mitochondria are visible as vertical and diagonal lines, respectively. (B) Quantification of percentage of moving mitochondria relative to the total number of mitochondria in patient sMNs compared to healthy controls. (C) Quantification of percentage of moving mitochondria relative to the total number of mitochondria at three different time points of sMN differentiation in *C9orf72* sMNs and healthy controls. Data for day 38 are reproduced from panel B. (D-F) Quantification of the percentage of moving mitochondria (marked with mitoDsRed2) relative to the total number of mitochondria in *C9orf72* sMNs and isogenic paired controls in which the hexanucleotide expansion has been excised. Two independent healthy controls are shown for comparison. Values are shown for mitochondria moving both anterogradely and retrogradely (D), anterogradely (E) or retrogradely (F). The difference in the magnitude of transport deficit between the experiment in B and C could be related to the different methods used to label mitochondria. Data are represented as mean ± S.D.; dots represent the number of neurites measured (see **Table S3** for total numbers of neurites analyzed in each experiment). Statistical significance was evaluated with Mann–Whitney test (B, C) or Kruskal-Wallis test with Dunn’s multiple comparison (D-F) (**** p<0.0001; *** p<0.001; ** p<0.01; *p<0.05; from N = 3 independent differentiations). In panel B a linear mixed model with random effect of subject and of experiment was also applied and confirmed significance.

To confirm a causal link between the *C9orf72* mutation and the transport deficit, we made use of three independent *C9orf72* iPSC lines and isogenic controls in which the repeat expansion had been corrected by CRISPR/Cas9 gene editing (49). In this set of experiments, mitochondria were visualized in iPSC-derived sMNs transduced with lentivirus expressing mitoDsRed2. There was a reduction in the frequency of axonal transport of mitochondria in the *C9orf72* sMNs compared to the paired mutation-corrected lines (Fig. 1D). The sparse lentivirus transduction of the sMNs allowed us to assess the directionality of mitochondrial transport in discrete neurons, revealing a significant inhibition of both retrograde and anterograde movements in the mutant cells (Fig. 1E, F). We conclude from these observations that transport of mitochondria in sMNs is impaired by the *C9orf72* repeat expansion.

### Arginine-rich DPRs disrupt microtubule-based transport in healthy motor neurons

As introduced above, there is compelling evidence that arginine-rich DPRs produced by *C9orf72* HREs contribute to pathophysiology (14,19–26,35,50). We therefore tested whether these molecules are sufficient to recapitulate the mitochondrial transport deficits observed in *C9orf72*-patient-derived motor neurons. We added peptides containing 20 repeats of PR or GR dipeptides (PR_20_ and GR_20_) to the culture medium of control iPSC-derived motor neurons. Consistent with previous observations (19, 26), these peptides were taken up by the motor neurons (**Fig. S7A-C**). We observed an inhibition of mitochondrial transport in motor neurons treated with arginine-rich DPRs at concentrations that did not trigger cell death (**Fig. S7D,** Fig. 2A, C-D and **Fig S8D-I**). A peptide with 30 PR repeats (PR_30_) produced a proportionately stronger impairment of mitochondrial transport (Fig. 2G and **Fig. S9A-D**), revealing a repeat-length-dependent effect. Delivery into cells of poly-GP (GP_20_), which has been shown to be non-toxic (21,36,37), did not affect motility of these organelles (Fig. 2A, B and **Fig. S8A-C**). Besides mitochondria, other cargos key to neuronal function are also trafficked along axons. These include RNA granules, which contain several proteins encoded by known ALS/FTD-associated genes (51, 52). We observed that transport of such granules (which can be marked with the SYTO-RNASelect dye ((51)) was reduced upon treatment with a non-lethal concentration of PR_20_ (Fig. 2E, F and **Fig. S8L-N**). We conclude that arginine-rich DPRs are sufficient to impair transport of cargos in human motor neurons.

**Fig. 2:**
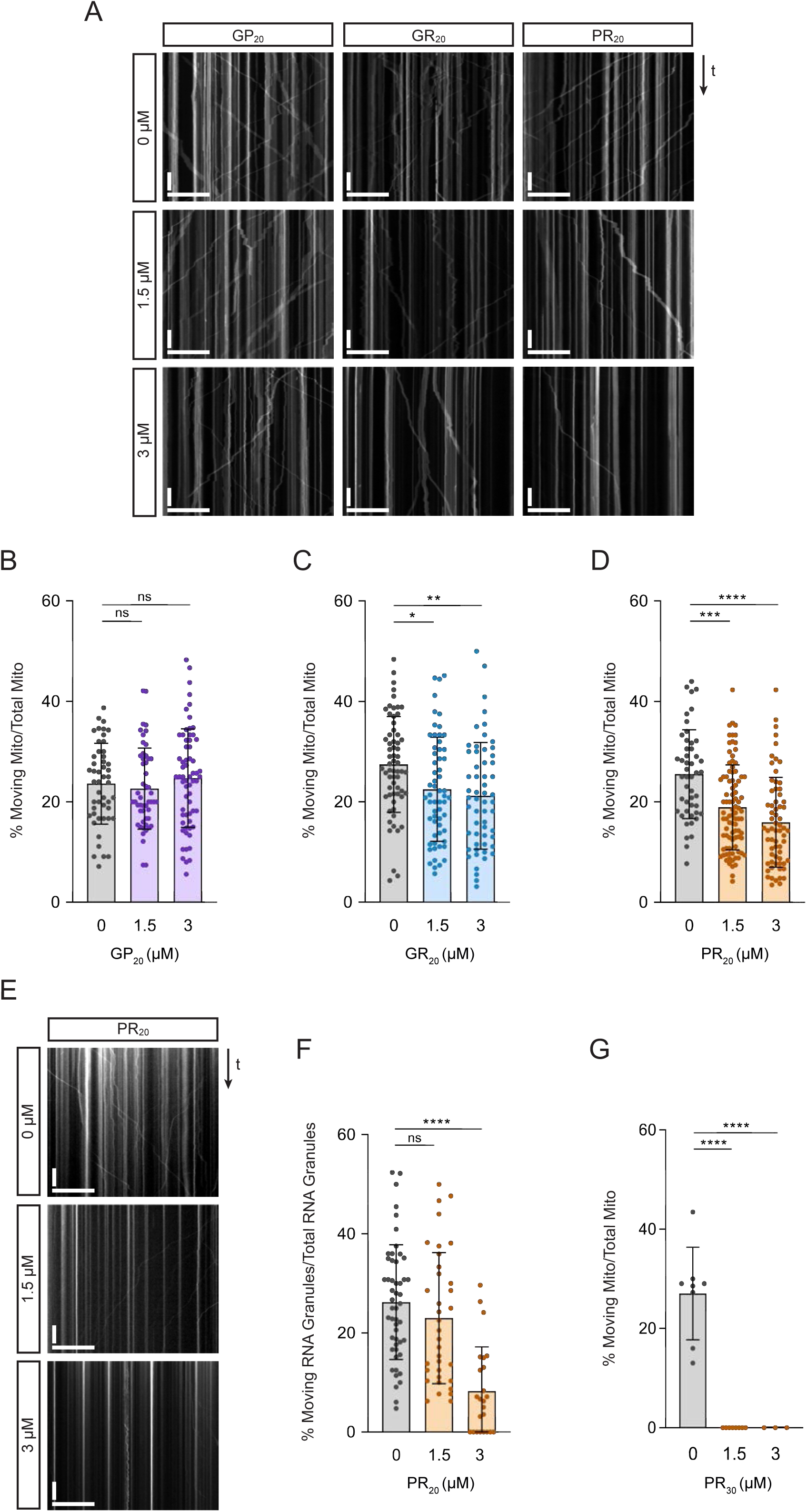
Arginine-rich DPRs impair transport of cargos in healthy sMNs. (A) Example kymographs of healthy sMNs after treatment with two different doses of synthetic GP20, GR20 and PR20 peptides. Y-axis, time; x-axis, distance; scale bars, 50 s and 30 µm. (B-D) Quantification of percentage of moving mitochondria relative to the total number of mitochondria upon treatment with GP20 (B), GR20 (C) or PR20 (D). (E) Example kymographs showing SYTO-RNA Select green-labeled stationary and moving RNA granules in healthy sMNs after treatment with two different doses of PR20. Y-axis, time; x-axis, distance; scale bars, 50 s and 30 µm. (F) Quantification of percentage of moving RNA granules relative to the total number of RNA granules after treatment with PR20. (G) Quantification of percentage of moving mitochondria relative to the total number of mitochondria after treatment with PR30. Data are represented as mean ± S.D.; dots represent the number of neurites measured (see **Table S3** for total numbers of neurites analyzed in each experiment). Statistical significance was evaluated with Kruskal-Wallis test with Dunn’s multiple comparison (B-D, F) or one-way ANOVA multiple comparisons test with Tukey’s correction (G) (**** p<0.0001; *** p<0.001; ** p<0.01; *p<0.05; ns, not significant; from N = 3 independent differentiations).

### Microtubule-based transport is perturbed by arginine-rich DPRs in a reconstituted *in vitro* **system**

From the experiments described above, we cannot conclude that arginine-rich DPRs inhibit the microtubule-based transport machinery directly. An alternative possibility is that perturbed axonal transport is a downstream consequence of other cellular processes being impaired. To test if these peptides can directly inhibit transport we used *in vitro* motility assays with purified motor complexes. Kinesin-1 and dynein were chosen for these experiments as they are the major motors for mitochondrial transport in axons (53–55), where they drive anterograde and retrograde movements, respectively. These motors also are also important for the transport of RNA granules in neuronal processes (56–58). To evaluate kinesin-1 motility we used a well characterized, constitutively active form of the major human isoform KIF5B tagged with GFP (59). Dynein motility assays were performed with the fluorescently-labelled human recombinant motor complex in the presence of the essential activators dynactin and BicD2N (60, 61). Experiments were performed with stabilized microtubules bound to a glass surface and concentrations of motor complexes low enough for visualization of single transport events by total internal reflection fluorescence (TIRF) microscopy.

We first assessed the effects of 20-repeat DPRs on motility of kinesin-1 by introducing them into the motility assay together with the motor complex (Fig. 3A). GR_20_ and PR_20_ significantly reduced the percentage of microtubule-associated kinesin-1 complexes that exhibited processive transport compared to the control in which no peptides were added (Fig. 3B). These DPRs also caused a significant decrease in the mean velocity of processive kinesin-1 movements, as well as an increase in the incidence of pausing events within or at the end of a run (Fig. 3B). GR_20_ and PR_20_ had the same consequences on the motility of dynein in the presence of dynactin and BicD2N, although the magnitude of these effects relative to the no-peptide control tended to be smaller than observed for kinesin-1 (Fig. 3C). In contrast, GP_20_ did not reduce the percentage of processive events or velocity of either motor, or increase their incidence of pausing (Fig. 3A-C).

**Fig. 3:**
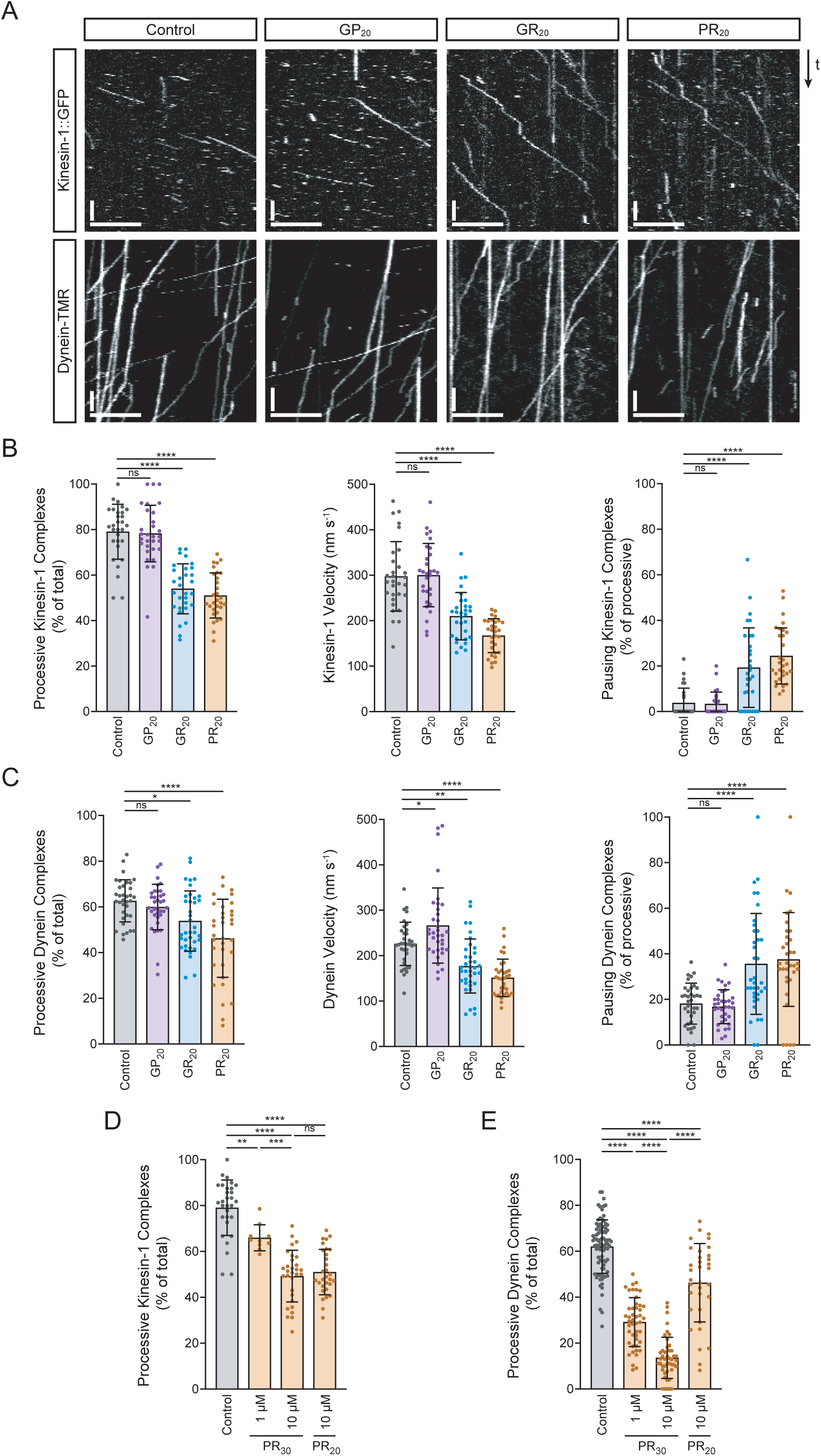
Arginine-rich DPRs perturb microtubule-based motility of kinesin-1 and dynein in a reconstituted *in vitro* system. (A) Kymographs showing examples of the behaviour of kinesin-1 (constitutively active 1-560 fragment tagged with GFP) and dynein (full complex with TMR-labelled heavy chain) on microtubules in the presence of 10 µM GP20, GR20, PR20 or no peptide (control). In all experiments in the study that assess dynein motility the activating co-factors dynactin and BicD2N were also present. In these kymographs, the microtubule plus-end is to the right. Y-axis, time; x-axis, distance; scale bars, 10 s and 5 µm. (B, C) Quantification of effects of 10 µM GP20, GR20 or PR20 on the percentage of microtubule-binding events that result in processive movement, velocity of processive movements, and the percentage of processive complexes that pause within or at the end of a run for kinesin-1 (B) and dynein (C). The values for dynein velocity are not statistically significantly different for GP20 vs control if the three outliers are omitted. (D, E) Quantification of effects of poly-PR chain length (10 µM PR20 or PR30) and concentration (1 µM or 10 µM PR30) on the percentage of microtubule-binding events of kinesin-1 (D) and dynein (E) that result in processive movement. In D and E, values for 10 µM PR20 and the control have been reproduced from B and C to facilitate comparison with other conditions. In B-E, means ± S.D. are shown; dots represent values for individual microtubules (see **Table S4** for total numbers of motor complexes analyzed in each experiment). Statistical significance was evaluated with a one-way ANOVA multiple comparisons test with Dunnett’s correction (B, C) or Tukey’s correction (D, E) (****p<0.0001; ***p<0.001; **p<0.01; *p<0.05; ns, not significant). See **Fig. S10** and **Fig. S11** for quantification of additional motile properties in these experiments.

Given the strong inhibition of mitochondrial transport in motor neurons by 30 repeats of PR (Fig. 2G), we also tested the effect of this peptide on both kinesin-1 and dynein complexes *in vitro*. Compared to controls lacking peptide, PR_30_ impaired processive movements of both kinesin-1 and dynein complexes in a dose-dependent manner (Fig. 3D and **Fig. S10A, B**). The effect of PR_30_ on dynein was proportionally stronger than observed for PR_20_ (Fig. 3E and **Fig. S10C**). In contrast, PR_30_ and PR_20_ impaired kinesin-1 motility to a similar extent (Fig. 3E and **Fig. S10D**). Upon further analysis, we found additional differences in the effect of arginine-rich DPRs on kinesin-1 and dynein. First, there were more overall binding events of kinesin-1 on microtubules in the presence of GR_20_, PR_20_ and PR_30_ compared to no peptide controls (**Fig. S11A** and **Fig. S10D**), whereas there were fewer such events for dynein (**Fig. S11B** and **Fig. S10C**). Second, whilst the arginine-rich DPRs prolonged the duration of processive movements for kinesin-1 complexes, the opposite was true for processive dynein complexes (**Fig. S11**).

These experiments demonstrate that arginine-rich DPRs perturb motility of both dynein and kinesin-1 in the absence of other potentially confounding cellular processes, although the effect of the peptides on the interactions of the two motors with microtubules are not equivalent.

### Arginine-rich DPRs bind microtubules *in vitro* and can impede motor movement

To shed light on how arginine-rich DPRs impair motor movement we monitored their localization in our *in vitro* motility assay. There was a clear interaction between fluorescently-labeled GR_20_, PR_20_ and PR_30_ and microtubules, with all three peptides binding along the length of the lattice as well as concentrating in puncta (Fig. 4A). The strongest overall binding was exhibited by PR_30_ (Fig. 4B), which also exhibited more frequent and more intense puncta than the 20-repeat DPRs (Fig. 4A). No microtubule association was observed for GP_20_ (Fig. 4A, B). Thus, only those DPRs that perturb motility of kinesin-1 and dynein are able to associate with microtubules.

**Fig. 4.**
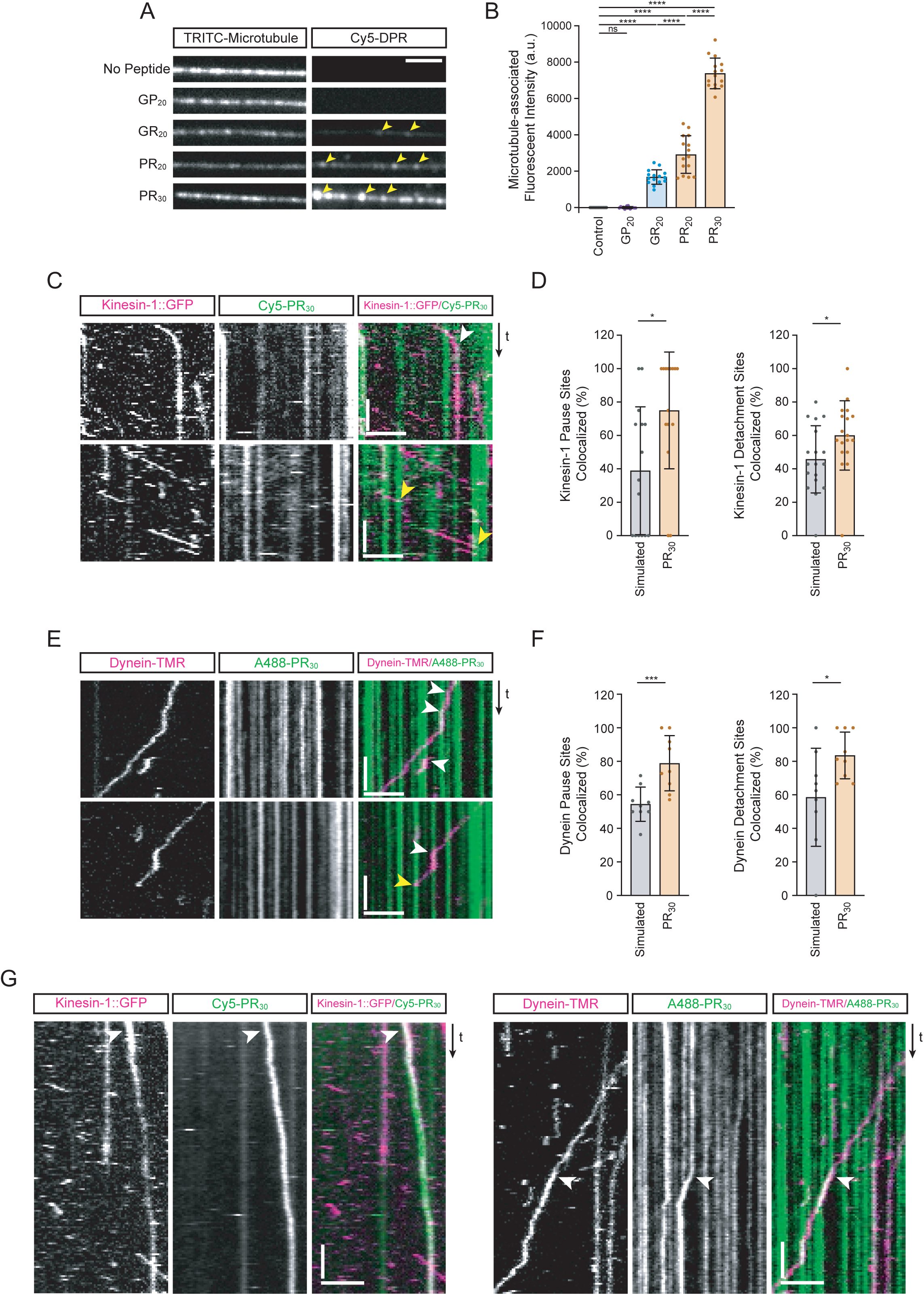
Arginine-rich DPRs bind microtubules *in vitro* and promote pausing or detachment of motors. (A) Example images of microtubules incubated with 10 µM GP20, GR20, PR20 or PR30. Arrowheads show examples of intense puncta of arginine-rich DPRs on microtubules. Scale bar, 2.5 µm. Uneven microtubule signal is due to stochastic incorporation of TRITC-tubulin into the lattice during polymerization. (B) Quantification of background-subtracted fluorescence intensity of DPR signals along the length of microtubules (chamber concentration of DPRs: 10 µM). (C-F) Analysis of localization of intense PR30 puncta with pausing or microtubule detachment events of motile kinesin-1 (C, D) and dynein (E, F) complexes. The chamber concentration of PR30 was reduced to 1 µM, which facilitated analysis of encounters of motors with regions of the microtubule with discrete DPR puncta. C and E, example of kymographs with white and yellow arrowheads showing, respectively, pausing and detachment of motors that localize with PR30 puncta. Y-axis, time; x-axis, distance; scale bars, 10 s and 2.5 µm. D and F, quantification of percentage of pausing or dissociation events of kinesin-1 (D) and dynein (F) localizing with intense PR30 puncta. Simulated controls were generated by overlaying motor trajectories with PR30 binding patterns from different microtubule regions (see Materials and Methods). (G, H) Kymographs showing examples (arrowheads) of the translocation of PR30 accumulations along microtubules by kinesin-1 (G) and dynein (H) (chamber concentration of PR30: 1 µM). Y-axis, time; x-axis, distance; scale bars, 10 s and 2.5 µm. In B, D and F, means ± S.D. are shown; dots represent values for individual kymographs. In these kymographs, the microtubule plus-end is to the right. See **Table S4** for total number of pausing and detachment events assessed in D and E. Statistical significance was evaluated with a one-way ANOVA multiple comparisons test with Tukey’s correction (B) or an unpaired t-test with Welch’s correction (D, F) (****p<0.0001; ***p<0.001; *p<0.05; ns, not significant).

These observations raised the possibility that microtubule-bound DPRs inhibit transport by impeding the movement of kinesin-1 and dynein along the track. Reducing the concentration of PR_30_ led to clearly resolvable puncta on microtubules (**Fig. S12**) that allowed us to investigate discrete interactions of DPRs and motor complexes on microtubules. We quantified the co-incidence of sites of pausing or microtubule detachment of kinesin-1 and dynein with the patches of intense PR_30_ accumulation and compared the results to control datasets generated with randomized patterns of PR_30_ foci at an equivalent density (Fig. 4C-F). Pause sites and detachment sites for both motors co-incided with a patch of PR_30_ significantly more often than expected by random chance (Fig. 4D, F). This result reveals that large accumulations of arginine-rich DPRs can act as obstacles to kinesin-1 and dynein complexes. Our observation that motor velocity along the microtubule was persistently reduced by poly-GR and poly-PR (Fig. 3A) suggests that the general accumulation of these DPRs along the lattice also affects motor motility.

We also detected instances of each type of motor complex transporting large foci of PR_30_ along microtubules (Fig. 4G; observed for 1.8% of processive kinesin-1 complexes (3/169) and 8.1% of processive dynein complexes (9/111)). Thus, in addition to interacting with microtubules, PR_30_ can associate with kinesin-1 and dynein-dynactin-BicD2N complexes in our reconstituted system. This observation led us to investigate whether arginine-rich DPRs can interact with microtubules and motors in the complex cellular milieu.

### Arginine-rich DPRs interact with microtubule and motor proteins in motor neuron extracts and *in vivo*

We first asked if PR_30_ interacts with tubulin and microtubule motors in mouse spinal cord lysate. Incubating the soluble fraction of the lysate with PR_30_ peptide, but not GP_30_, lead to the formation of large biomolecular condensates that could be pelleted by gentle centrifugation and analyzed by mass spectrometry (36) (Fig. 5A, B). The spinal cord interactome of PR_30_ was strongly enriched for similar gene ontology categories (e.g., RNA granule, translation, proteasome, protein folding) as observed in our previous interactome study using cancer cell extracts (36) (Fig. 5C, **Fig. S13A, B** and **Table S5**). We also identified a strong enrichment of terms related to intracellular transport (Fig. 5C, D). Proteins in this category included those involved in nucleocytoplasmic transport (**Table S5**), a process we and others previously implicated in *C9orf72* ALS/FTD (19,21,22,25,35), and microtubule-based transport (Fig. 5C). Consistent with the interactions of arginine-DPRs observed in our *in vitro* motility assays, several isotypes of the α- and β-tubulin subunits of microtubules were precipitated with PR_30_ (Fig. 5D and **Table S5**), as were the three different kinesin-1 isoforms (KIF5A, KIF5B and KIF5C) and components of the dynein and dynactin complexes (Fig. 5D and **Table S5**). We also detected other classes of kinesins and components of intraflagellar transport dynein (cytoplasmic dynein-2) in the precipitate (Fig. 5D and **Table S5**). These data reveal that PR_30_ can associate in spinal cord lysate with multiple proteins involved in microtubule-based transport (Fig. 5D and **Table S5**). Several of these factors are mutated in ALS or other neuromuscular disorders (Fig. 5D**, Fig. S13A, B** and **Table S5**).

**Fig. 5:**
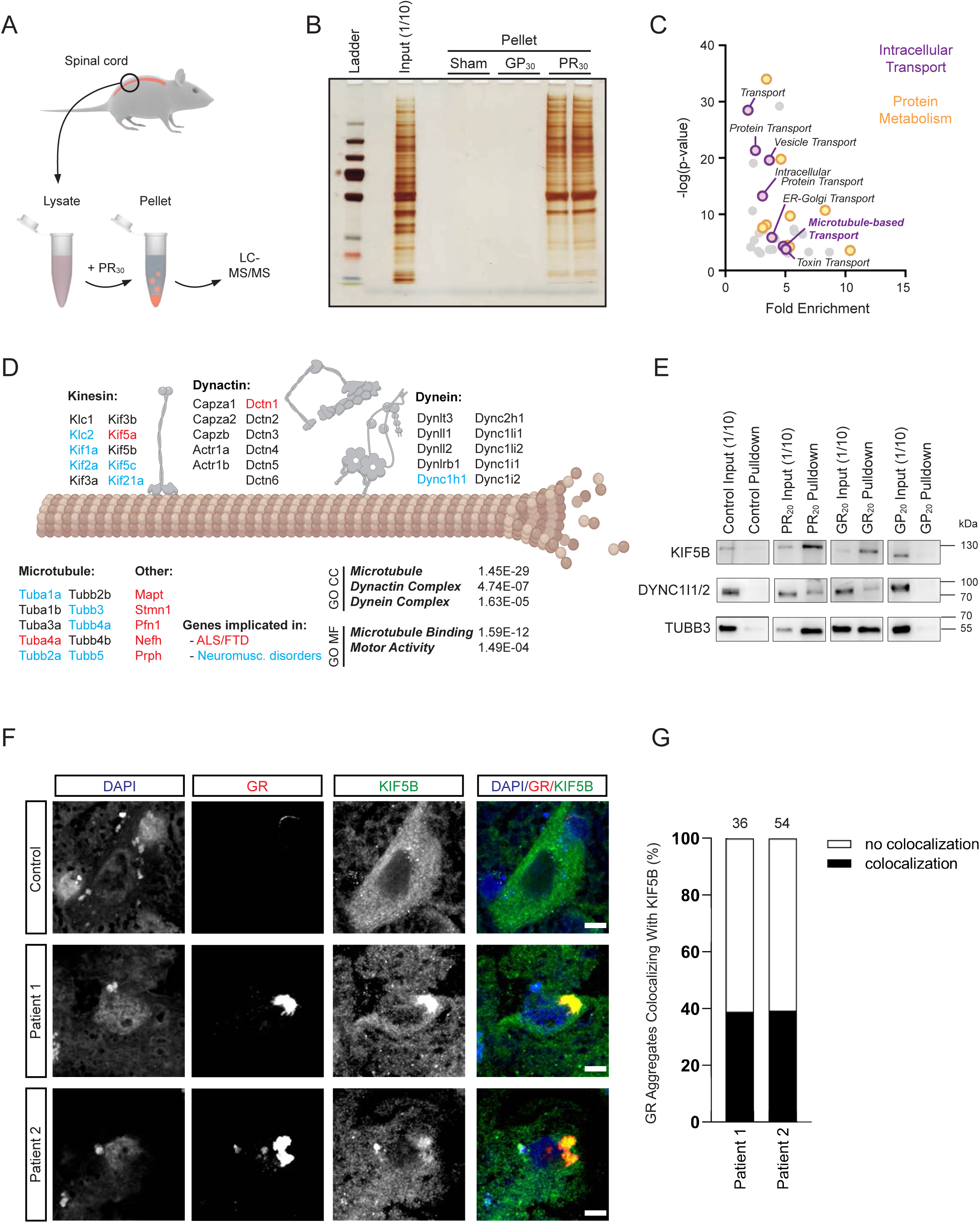
Arginine-rich DPRs interact with axonal transport machinery. (A) Overview of mass spectrometry experiment. Soluble mouse spinal cord lysate was incubated with PR30 to induce protein phase separation. The resulting coacervate was precipitated by centrifugation and analyzed by mass spectrometry. (B) Silver stain of pellet fractions show that PR30, but not GP30 induces protein phase separation. (C) Proteins in the PR30 interactome (n = 1811) were enriched for GO categories centered around protein metabolism (including factors involved in translation, folding and degradation) and intracellular transport. (D) Illustration of components of the axonal transport machinery present in the PR30 interactome (GO CC: GO analysis cellular compartment; GO MF: GO analysis molecular function). Proteins associated with ALS/FTD or other neuromuscular disorders are highlighted in red and blue, respectively. (E) Immunoblots confirming interactions of KIF5B, dynein and tubulin with PR20 and GR20, but not GP20, in human sMNs lysates (for each protein, images are crops from the same gel, see **Fig. S13C-E** for the uncropped gels). In (B) and (E) the input was diluted 1:10 before gel loading. (F) Immunofluorescence of frontal cortex from two *C9orf72* patients showing examples of enrichment of KIF5B in poly-GR aggregates. ALS patients without a *C9orf72* expansion were used as controls. Scale bar, 5 µm. (G) Quantification of co-localization of poly-GR inclusions and KIF5B foci (numbers of poly-GR inclusions analyzed for each patient are shown above the bars).

The association of arginine-rich DPRs with kinesin-1, dynein and microtubules was further supported by treating control iPSC-derived sMNs with HA-labelled PR_20_, GR_20_ and GP_20_ peptides and subjecting their lysates to immunoprecipitation using anti-HA beads. Robust signals for KIF5B, tubulin and dynein were observed in immunoblots of the material precipitated with PR_20_ and GR_20_, but not the material precipitated with GP_20_ (Fig. 5E and **Fig. SC-E**).

We next investigated interactions of DPRs with microtubules and motors within intact sMNs using fluorescence microscopy. The broad distribution of microtubules, KIF5B and dynein within cell bodies and along neuronal processes prevented meaningful analysis of interactions with exogenous DPRs at these sites by conventional co-localization analysis. However, co-localization of signals from poly-PR and tubulin could be detected in growth cones of developing sMNs, in which microtubules are spaced apart (**Fig. S14A**). Furthermore, duolink proximity ligation assays (62, 63) supported the interaction between PR_20_ and KIF5B in both the cell body and processes of sMNs (**Fig. S14B**). We also investigated the association of KIF5B with poly-GR in brain sections from two patients with *C9orf72* ALS/FTD. Approximately 40% of the cytoplasmic poly-GR inclusions in the sections had enrichment of KIF5B, supporting the association of endogenously expressed arginine-rich DPRs with this motor (Fig. 5F, G). Collectively, these observations strengthen the evidence for physical association of arginine-rich DPRs with the axonal transport machinery.

### Altering levels of kinesin-1 strongly modifies defects caused by expression of arginine-rich DPRs in *Drosophila*

The experiments described above provide evidence that arginine-rich DPRs impair transport directly by interacting with microtubules and motors. However, these data do not indicate whether this activity contributes to the pathomechanism of arginine-rich DPRs *in vivo*. To address this question, we manipulated the microtubule-based transport machinery in previously described *Drosophila* models for toxicity of these peptides (20). We took advantage of a transgene containing the entire *Kinesin-1 heavy chain* (*Khc*) locus (*P[Khc^+^]*) that increases Khc protein levels by approximately two-fold and stimulates axonal transport of mitochondria in neurons of aged adult flies (64, 65). Both anterograde and retrograde movements of these organelles are more frequent in these animals, consistent with tight coupling of dynein and kinesin-1 motor activities in many cell types (66–68).

We first used flies overexpressing 36 repeats of GR under control of the GMR-GAL4 driver. It has previously been reported that expressing GR_36_ in this manner results in a high incidence of lethality before adulthood (20). We also observed substantial lethality in this condition, with ∼40% of the expected number of adult flies recovered (Fig. 6A). Remarkably, *P[Khc^+^]* almost completely suppressed the lethality caused by GR_36_ expression (Fig. 6A). Consistent with this result, inactivating one copy of the endogenous gene encoding Khc significantly enhanced the lethal phenotype compared to flies with wild-type levels of Khc (Fig. 6A).

**Fig. 6:**
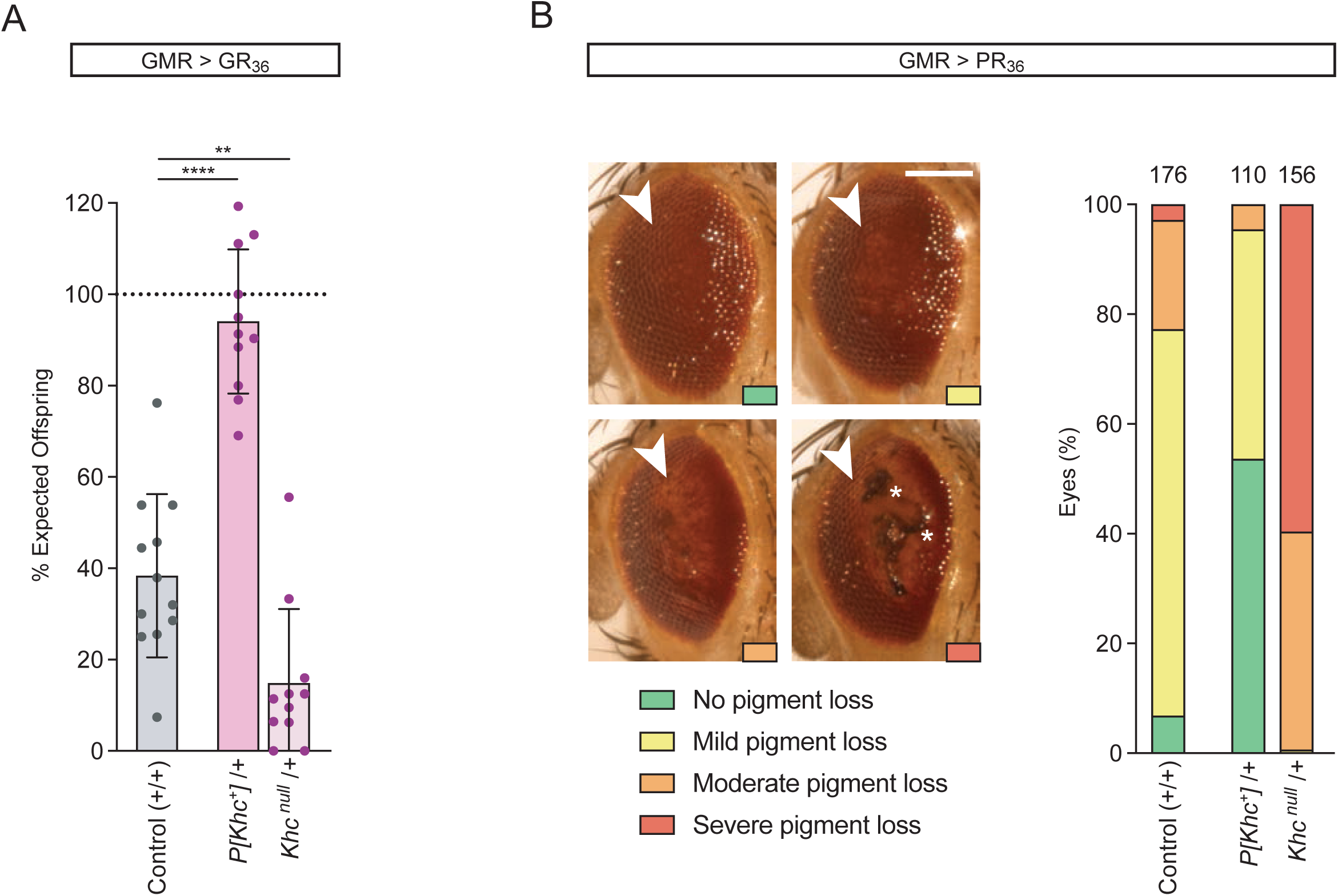
Altering gene dosage of kinesin-1 strongly modifies lethality and eye defects caused by poly-GR or poly-PR expression in *Drosophila*. (A) Quantification of effect of Kinesin-1 heavy chain (Khc) gene dosage on the lethal phenotype caused by GR36 overexpression. Adult flies were scored 4-6 days after eclosion. Khc was overexpressed with a transgene containing the genomic locus (*P[Khc^+^]*). Data are expressed as the percentage of offspring of this genotype expected when there is no lethality. Means ± S.D. are shown; dots represent values for each egg lay. Statistical significance was evaluated with a one-way ANOVA multiple comparisons test with Dunnett’s correction (****p<0.0001; **p<0.01). (B) Effects of Khc gene dosage on the eye defects caused by PR36 overexpression. Left, representative images of phenotypic categories. Note that all fly eyes retained the characteristic rough eye phenotype of PR36 over-expression, regardless of Khc gene dosage. Arrowheads show regions with pigment loss. Severe pigment loss was often accompanied by the appearance of necrotic regions (asterisks). Scale bar, 100 µm. Right, quantification of eye phenotypes (total number of eyes analyzed shown above bars). Data are compiled from several crosses per genotype; see **Fig. S15** for data from individual crosses and statistical evaluations of phenotypic changes.

Expression of arginine-rich DPRs also results in a range of abnormalities in the adult eye, including disruption to the external lattice, pigment loss and necrosis (20, 25). *P[Khc^+^]* also suppressed eye abnormalities caused by GR_36_ expression compared to a control with wild-type levels of the motor (**Fig. S15A**). The strong lethality of GR_36_-expressing *Khc* heterozygous mutants meant there were not enough animals for meaningful analysis of eye phenotypes. To more comprehensively investigate the link between motor levels and DPR-induced eye defects, we used a PR_36_-expressing fly stock that causes eye abnormalities without compromising development to adulthood (20). Consistent with the genetic interactions with GR_36_, the severity of eye defects caused by PR_36_ was suppressed by *P[Khc+]* and enhanced by reducing *Khc* gene dosage (Fig. 6B and **Fig. S15B, C**). In control experiments, we determined that in the absence of GR_36_ or PR_36_ the changes in *Khc* gene dosage had no effect on survival or eye morphology (**Table S6**).

Collectively, these data reveal that increasing the amount of the kinesin-1 motor suppresses the defects caused by overexpression of poly-GR and poly-PR, which further supports a functional link between the microtubule-based transport machinery and the pathomechanism of arginine-rich DPRs.

## DISCUSSION

Axonal transport of organelles and macromolecules by microtubule motors is crucial for neuronal homeostasis and survival. This process facilitates communication between synaptic terminals and the cell body and allows local demands for the functions of organelles and macromolecules to be met (44). Deficits in microtubule-based transport in axons are linked with several age-related neurodegenerative diseases including Alzheimer’s, Huntington’s disease and hereditary spastic paraplegia (43), as well as with ALS caused by mutations in *SOD1*, *TARDBP* and *FUS* (44– 48,51). However, whether impaired axonal transport is a cause or consequence of neuronal dysfunction in these contexts remains controversial. The notion that defective intracellular trafficking can directly trigger ALS has been strengthened by the discovery of disease-causing mutations in several genes encoding components of the microtubule-based transport machinery, namely TUBA4A, DCTN1 and KIF5A (40–42). These mutations have been reported to disturb the dynamics of microtubules or the trafficking of cargos along them (40,41,69–73).

Despite evidence of defective axonal transport in other forms of ALS, the literature on *C9*-ALS/FTD has focused on other potential disease mechanisms (74). Here, we report that the disease-causing HREs in the *C9orf72* gene impair transport in neurites of human iPSC-derived sMNs. These data indicate that deficits in intracellular trafficking of cargos are a common feature of ALS pathogenesis.

Although both loss- and gain-of-function disease mechanisms have been proposed in *C9*-ALS/FTD (74), the transport impairment in the patient-derived motor neurons was not associated with a reduction in overall *C9orf72* mRNA levels. However, we found that administration of the arginine-rich DPRs that are produced from *C9orf72* HREs inhibits transport of mitochondria and RNA granules in sMNs derived from healthy individuals. These observations, which are in line with the ability of PR_36_ to increase the stationary fraction of mitochondria in a *Drosophila* model (46), suggest a primarily gain-of-function mechanism in human neurons involving inhibition of axonal transport by poly-PR and poly-GR. The relevance of perturbed microtubule-based transport to the pathomechanism of arginine-rich DPRs is supported by our finding that changes in kinesin-1 gene dosage strongly modify the toxic effects of arginine-rich DPRs in *Drosophila*. Most strikingly, we found that boosting levels of kinesin-1, which we previously showed increases the frequency of anterograde and retrograde axonal transport in adult flies (65), almost completely suppresses the lethality caused by GR_36_ expression.

Our *in vitro* motility assays reveal that poly-GR and poly-PR can directly perturb microtubule-based transport of kinesin-1 and dynein, which are key motors in the trafficking of many cargos in neurons, including mitochondria (53–55) and RNA granules (56–58). These peptides have a range of effects on motor behavior, including reducing the fraction of microtubule-associated complexes that are processive, as well as slowing the movement of processive complexes and increasing their incidence of pausing. We show that arginine-rich DPRs can associate with microtubules *in vitro* suggesting that they directly interfere with motor translocation along the track. Consistent with this notion, large puncta of microtubule-associated PR_30_ can act as obstacles for kinesin-1 and dynein, stimulating motor pausing or detachment. This effect is reminiscent of the ability of the MAPT (tau) protein to partition into patches on microtubules and obstruct the motility of kinesin-1 and dynein complexes *in vitro* (75–79). Mutations in both *MAPT* and *C9orf72* have been shown to cause FTD (80) and it was recently proposed that this reflects a shared inhibitory effect on nucleocytoplasmic transport (81). The ability of both arginine-rich DPRs and MAPT to impede the motors responsible for transport in the cytoplasm offers another possible explanation for the overlapping features of *C9*-ALS/FTD and MAPT-FTD.

We also detect physical association of arginine-rich DPRs with kinesin-1 and components of the dynein-dynactin transport complex, which could also contribute to the inhibition of transport. Indeed, the reduced velocity and increased pausing of both types of motor in the presence of arginine-rich DPRs *in vitro* is compatible with their retardation by transient interactions with microtubule-associated peptides.

Intriguingly, not all properties of kinesin-1 and dynein are affected equivalently by arginine-rich DPRs. While the peptides increase the frequency of kinesin-1 binding to microtubules and the duration of its processive runs, the opposite is true for dynein. These observations indicate that the overall interactions of kinesin-1 and dynein complexes with microtubules are increased and decreased, respectively, by arginine-rich DPRs. Whilst the mechanistic basis of this differential effect warrants further investigation, our current results highlight how either strengthening or weakening interactions with microtubules can prevent efficient translocation of motors along the track.

We found that transport of both mitochondria and RNA granules in sMNs is perturbed by arginine-rich DPRs. Given the widespread roles of kinesin-1 and dynein in intracellular transport (82, 83), as well as our discovery of other classes of microtubule motors in the PR_30_ interactome from sMNs, it is likely that the trafficking of several cargos is disrupted by these peptides. Whether altered translocation of a specific cargo type is of particular significance for *C9-*ALS/FTD remains to be resolved. However, the observation that ALS/FTD-associated mutations in the RNA-binding protein TDP-43 alter its bidirectional transport along microtubules in axons (51) suggests that RNA trafficking abnormalities contribute to disease pathology. Consistent with this hypothesis, TDP-43 has been detected, like KIF5B in this study, in endogenous GR inclusions in *C9*-ALS/FTD post-mortem tissue (84).

Whilst our results support a direct inhibitory effect of arginine-rich DPRs on axonal transport, we cannot rule out that other processes also contribute to the trafficking defects we observe in *C9*-ALS/FTD neurons. In addition to defective nucleocytoplasmic transport, altered RNA splicing, translation, and membrane-less organelle dynamics, as well as elevated mitochondrial and ER stress, have been linked to the disease (14,21,22,25,26,28–33,35–38). It is possible that one or more of these processes exacerbates the transport defects associated with binding of arginine-rich DPRs to the axonal transport machinery. Nonetheless, our data indicate that manipulating microtubule-based transport could be an effective strategy to hinder the progression of *C9*-ALS/FTD, possibly in conjunction with therapies that target other processes affected by the HREs.

## MATERIAL AND METHODS

### Cell culture and motor neuron differentiation

*C9orf72* iPSC lines were generated from dermal fibroblasts from ALS (C9-2B and C9-3B) and ALS/FTD (C9-1B and C9-4B) patients carrying the G_4_C_2_ repeat expansion in the *C9orf72* gene (**Table S1**). Controls were generated from unrelated healthy individuals. Con 1B and Con 2B were purchased from Takara Bio Inc (ChiPSC-6b, P11031) and Sigma (iPSC Epithelial-1, IPSC0028), respectively. Con 3B, C9-1B, C9-2B, C9-3B and C9-4B were generated using the CytoTune™-iPS 2.0 Sendai Reprogramming Kit (ThermoFisher Scientific, A16517). qRT-PCR for Sendai virus clearance was performed and did not detect residual Sendai virus RNA in any of the tested iPSC lines. iPSCs were kept in in Essential 8™ flex medium (A28583-01, Gibco^TM^) with 1000 U/ml penicillin–streptomycin. Colonies were passaged every week with 0.5 mM EDTA (15575-020, Invitrogen) diluted in Dulbecco’s phosphate-buffered saline (DPBS) and plated on GeltrexR LDEV-Free, hESC-Qualified, Reduced Growth Factor Basement Membrane Matrix (A1413302, Gibco^TM^). The absence of mycoplasma contamination was routinely confirmed by PCR. Controls (Con-1B, Con-2B and Con-3B) and *C9orf72* (C9-1B, C9-2B, C9-3B and C9-4B) iPSCs were differentiated into spinal motor neurons as previously described (45, 85). The controls (Con-1 and Con-2), the isogenic controls (C9-1Δ, C9-2Δ, C9-3Δ) and the *C9orf72* counterparts (C9-1, C9-2 and C9-3) were obtained and differentiated as reported in Selvaraj et al., 2018 (49).

### Immunocytochemistry

Coverslips were incubated for 20 min at room temperature in phosphate-buffered saline (PBS) with 4% paraformaldehyde and washed three times with PBS. Cells were incubated for 1 h at room temperature in PBS containing 0.1% Triton X-100 (Acros Organics) and 5% donkey (Sigma) or goat (Dako) serum. Cells were incubated with primary antibodies diluted in PBS containing 0.1% Triton X-100 and 2% serum overnight at 4 °C. Cells were rinsed three times in PBS before the secondary antibody was added for 1 h at room temperature. See **Table S7** for details of primary and secondary antibodies. After washing with PBS, the coverslips were incubated for 20 min in PBS/DAPI (NucBlue™ Live ReadyProbes™ Reagent, Invitrogen, R37605). Coverslips were rinsed three times with PBS before mounting in ProLong™ Gold antifade reagent (Invitrogen, P36934). Confocal images were obtained using a Leica SP8 DMI8 confocal microscope. Images were analyzed, formatted and quantified with ImageJ software.

### Pluripotency characterization

iPSCs cells were subjected to spontaneous differentiation mediated by the formation of embryoid bodies (EBs), and subsequently analyzed for the three-lineage differentiation using the ScoreCard methodology (86) (ThermoFisher Scientific). On the day of EB formation, 60-80% confluent iPSCs were washed with PBS and incubated with 0.5 mM EDTA (15575-020, Invitrogen) for 1-3 min to dissociate the colonies. Cells were harvested in Essential 8™ flex medium (A28583-01, Gibco^TM^) and counted before gently spinning down at 300 x *g* for 5 min. Cells were plated in a 24 well Corning® Ultra-Low Attachment Surface plates (Corning, 3473) at the density of 1.5 × 10^6^ cells/well. The plates were incubated overnight at 37°C. The following day, half of the Essential 8™ flex medium was replaced with Essential 6™ medium (A15164-01, Gibco^TM^). For the spontaneous differentiation, the EBs were kept for 14 days in the E6 medium, which was changed every 2 days. After 14 days in culture, EBs were collected for RNA extraction with the GenElute Mammalian Total RNA kit (Sigma, RTN350). cDNA synthesis was performed with Superscript^TM^ III (ThermoFisher Scientific, 11752-050) and qPCR was performed according to the manufacturer’s protocol with TaqMan hPSC Scorecard assay (ThermoFisher Scientific, A15871). Data were analyzed with the online Scorecard software (ThermoFisher Scientific).

### Real-time PCR

Total RNA extraction was performed with the GenElute Mammalian Total RNA kit (Sigma). cDNA was synthesized from 1 μg of total RNA using SuperScript® III First-Strand Synthesis SuperMix for qRT-PCR (Invitrogen) according to the manufacturer’s instructions. The qPCR for Sendai virus detection was performed using TaqMan Gene Expression Assays (Life Technologies). Quantitative RT-PCR was performed using SYBR® Green PCR Master Mix (Applied Biosystems™) on the 7500 Step OnePlus™ Real-Time PCR System (Applied Biosystems™). Relative gene expression was determined by the 2−ΔΔct method with normalization to *GAPDH* mRNA.

### Repeat primed PCR

The *C9orf72* repeat expansion mutation was confirmed in both iPSCs and MNs by repeat primed PCR (RP-PCR). First, the G_4_C_2_ hexanucleotide repeat was amplified using the forward primer *C9orf72*-PCR-F (6-FAM) CAAGGAGGGAAACAACCGCAGCC and the reverse primer *C9orf72*-PCR-R GCAGGCACCGCAACCGCAG. The PCR reaction mix was prepared in a 25 μl solution consisting of: 100 ng of genomic DNA, 10X PCR amplification buffer, 0.5 μl of Taq DNA polymerase (Roche, 1435094), 2.5 μM forward primer, 2.5 μM reverse primer, 2 mM deoxyribonucleotide triphosphate (dNTP) (Amersham Biosciences, 272094-02), 50 mM MgSO_4_ (Invitrogen), 6 μl of PCR enhancer solution (Invitrogen, 11495-017) and dH_2_O to the final volume. After a 5-min incubation at 94 °C, the reactions were subjected to 28 cycles of 95 °C for 30 s, 55 °C for 30 s, 68 °C for 60 s, followed by a 5 min extension at 68 °C. A forward primer *C9orf72*_F_FAM_bis (6-FAM)-AGTCGCTAGAGGCGAAAGC, a reverse primer *C9orf72*_R1_bis TACGCATCCCAGTTTGAGACGGGGGCCGGGGCCGGGGCCGGGG, and a second reverse primer *C9orf72*_R2_bis TACGCATCCCAGTTTGAGACG were used for the amplification of the G_4_C_2_ hexanucleotide repeat. The RP-PCR assay was performed in a reaction volume of 25 μL containing: 100 ng genomic DNA, 10X expand long template buffer 2 (Roche, 11681842001), expand long template enzyme mix (Roche, 11681842001), 2.5 μM primer F, 2.5 μM primer R1, 2.5 μM primer R2, 20 mM dNTPs (Amersham Biosciences), 5 mM betaine (Sigma, B0300-1VL) and water to the final volume. After a 10 min incubation at 98 °C, the reactions were subjected to 10 cycles of denaturation at 97 °C for 35 s, 53 °C for 2 min, 68 °C for 2 min, followed by 25 cycles (+20 s/cycle) of 97 °C for 35 s, 53 °C for 2 min and 68 °C for 2 min. This was followed by a 10 min extension at 68 °C. PCR products were incubated at 95 °C for 3 min and cooled on ice followed by 5 μl of each PCR product being added to 18 μL of formamide (Sigma) and 2 μL of Genescan 500 Rox Size Standard (Applied Biosystems). Fragment length analysis was performed following electrophoresis on an automatic sequencer (ABI3730 DNA Analyzer; Applied Biosystems), using GeneMapper software version 4.0 (Applied Biosystems). A characteristic stutter amplification pattern on the electropherogram was considered diagnostic of a pathogenic repeat expansion.

### Southern blotting

Southern blots were performed as described previously (87), briefly genomic DNA was extracted using the Wizard Genomic DNA Purification Kit (Promega) and digested with the XbaI restriction enzyme (Promega). After an electrophoresis of six hours at 100 V on a 0.8% agarose gel, DNA was transferred to a positively charged nylon membrane (Roche) and then crosslinked by UV irradiation. Hybridization was performed using a digoxigenin (DIG)-labelled probe (Roche). The probe was detected with an anti-DIG antibody (Roche; 1:10,000) and visualized using CDP-star substrate (Roche) and autoradiography film.

### Electrophysiology

Whole-cell patch-clamp recordings from iPSC-derived motor neurons were performed at room temperature in artificial cerebrospinal fluid (aCSF) consisting of: 140 mM NaCl, 5 mM KCl, 2 mM CaCl_2_, 2 mM MgCl_2_, 10 mM HEPES and 12 mM glucose (pH 7.4; ∼300 mOsm). Borosilicate glass patch pipettes were pulled by a vertical PIP6 micropipette puller (HEKA Elektronik, Lambrecht/Pfalz, Germany) and filled with an internal solution containing: 120 mM K-gluconate, 20 mM KCl, 1 mM MgCl_2_, 10 mM HEPES, 0.2 mM EGTA, 0.3 mM Na-GTP, 5 mM NaCl, 4 mM Mg-ATP (pH 7.3; ∼290 mOsm). Motor neurons were visualized with an inverted Olympus IX73 microscope equipped with a 40X objective (0.60 NA, Ph2). Action potentials were recorded in current-clamp mode. Signals were acquired, filtered (at 2.8 kHz) and digitized (at 20 kHz) using an EPC10 USB amplifier and PatchMaster software (HEKA Elektronik). The liquid junction potential of 13.8 mV was corrected off-line. Action potentials were detected using Stimfit software.

### Digital droplet polymerase chain reaction

The QX200 Droplet Digital PCR (ddPCR) system (Bio-Rad) was used to provide absolute measurements of *C9orf72* mRNA, as well as each of the V1, V2, V3 transcript variants. Total RNA was extracted with the RNeasy Mini Kit*®* following the manufacturer’s instructions. The cDNA was obtained using the SuperScript*®* III First-Strand Synthesis System for RT-PCR kit according to the manufacturer’s instructions. Digital droplet PCR was performed following Bio-Rad guidelines (Bio-Rad). Briefly, each ddPCR reaction mix was prepared in a 22 µl solution consisted of 2X ddPCR Supermix for probes (without dUTPs), 20x FAM Taqman probe, 20x Hex Taqman probe, 4 µl of diluted cDNA (1/10), and 4.8 µl of nuclease-free water (#R0581). Droplets were generated in a QX200 Droplet Generator and transferred to a standard 96-well PCR plate that was heat-sealed using foil sheets (Pierceable Foil Heat Seal, Bio-Rad, 1814040) and the PX1 PCR plate sealer. The droplets were amplified using the Bio-Rad T100 thermal cycler starting with 10 min enzyme activation at 95 °C, followed by 40 cycles of 95 °C, 30 s and 60 °C, 1 min and a final hold at 98 °C for 10 min (2 °C/s ramp rate). The fluorescence of each thermal cycled droplet was measured using the QX200 Droplet Reader (Bio-Rad) and the results analyzed with QuantaSoft software (Bio-Rad) after threshold setting on fluorescence of negative controls. The number of positive and negative droplets was used to calculate the concentration (cDNA copies/μl of the final 1x ddPCR reaction) of the targets and their Poisson-based 95% confidence intervals. A reference gene *SCLY* was included in order to increase the accuracy of the relative gene expression. Graphs show relative expression levels. The V1: 6-FAM-TAA TGT GAC-ZEN-AGT TGG AAT GC-IBFQ, V2: 6-FAM-CGG AGC ATT-ZEN-GG ATA ATG TGA CAG TTG G-IBFQ, V3: 6-FAM-TAA TGT GAC-ZEN-AGT TGG AAT GC-IBFQ and Vall: 6-FAM-ACA GAG AGA-ZEN-ATG GAA GAT CAG GGTC AGA-IBFQ probes were used in this experiment (6-FAM = 6-carboxyfluorescein, ZEN = internal quencher, IBFQ = Iowa Black dark fluorescence quencher).

### Enzyme-linked immunosorbent assay (ELISA)

Poly-GP and poly-GA levels in control and *C9orf72* derived sMNs lysates were measured using a sandwich immunoassay that utilizes Meso Scale Discovery (MSD) electrochemiluminescence detection technology. Lysates were diluted in assay diluent (1% Blocker-A/PBST) and equal amounts of protein for all samples were tested for poly-GP. The assay employs an affinity purified goat polyclonal antibody to poly-GP (Biogen-3746 1.0 µg/mL) as capture, and mouse monoclonal poly-GP antibody (Biogen 2P8H9.1.1, 0.5 µg/mL) along with a SULFO-tag anti-mouse secondary antibody (1.0 µg/ml), to detect captured poly-GP. Similarly, lysates were diluted in assay diluent (1% Blocker-A/PBST) and equal amounts of protein for all samples were tested for poly-GA using an affinity purified goat polyclonal antibody to poly-GA (Biogen-3682, 1.0 µg/mL) as capture, and mouse monoclonal to poly-GA antibody (Biogen 2P36E2.1.1, 1.0 µg/mL) along with a SULFO-tag anti-mouse secondary antibody (1.0 µg/ml). Each sample was tested in duplicate wells, sample volume permitting, using 50 µL per well. Each assay plate contained the same control samples (serial dilutions of iPSC motor neuron cell lysates derived from a *C9orf72* mutation carrier and a non-carrier, and serial dilutions of synthetic GP_8_ or GA_8_. Response values reported correspond to intensity of emitted light upon electrochemical stimulation of the assay plate using the MSD QUICKPLEX SQ120, from which the background response in wells containing only assay buffer was subtracted.

### Cell viability assay

Cell viability after DPR treatment was measured using DRAQ7™ (Abcam, ab109202) far-red fluorescent DNA dye. DRAQ7 is a cell impermeable dye that only stains the nuclei in cells with a compromised cell membrane, i.e. that are in the process of dying. DRAQ7 was added to each well at a concentration of 3 µM together with Hoechst 33342 (1:1000 Thermo Fisher Scientific, H1399) and incubated for 20 min at 37 °C. Cells were then washed once with DMEMF12 and fixed with 4% paraformaldehyde in PBS for 20 min. For this experiment, motor neurons were cultured in a CellCarrier 96-well plate (Perkin Elmer, UK). Motor neurons were treated with the respective synthetic peptides (PR_20_, GR_20_, GP_20_; Pepscan) and the analysis was performed 24 h after treatment. Untreated cells were used as the control group for PR_20_ and GR_20_. To allow the uncharged peptide, GP_20_, to pass the cellular membrane, cells were treated with Streptolysin O (SLO, 15 ng/ml) previously diluted in HBSS (without Ca^2+^), Gibco, 14175-053) containing 30 mM HEPES (Sigma, H3375) for 15 min, 37 °C. In this case, cells treated with SLO without GP_20_ peptide were used as untreated controls. Images were acquired using the Operetta CLS High Content Screening (HCS) System (Perkin Elmer, UK) using a 20X objective lens (0.4 NA dry, correction collar). Each experimental condition was assayed in at least three wells for each of at least two independent experiments. 77 fields of view were acquired per well. Images were analyzed using Harmony software (PhenoLOGIC, PerkinElmer).

### Visualization and analysis of cargo transport in neurons

Motor neuron cultures were stained with MitoTracker^TM^ Red (50 nM, Invitrogen, M22425) or SYTO RNASelect^TM^ Green fluorescent (500 nM, Invitrogen, S32703) for 20 min, at 37 °C, according to the manufacturer’s instructions. Control sMNs were treated with the respective synthetic peptides for 24 h prior analysis of cargo transport, as described for the cell viability assay. Live cell imaging was performed using an inverted Zeiss Axiovert 200 M microscope (Carl Zeiss) with a 40X water immersion lens (1.2W, C-Apochromat). During the recording, the cells were perfused with HEPES solution (10mM HEPES, 148 mM NaCl, 5 mM KCl, 0.1 mM MgCl_2_, 10 mM glucose, 2 mM CaCl_2_) and maintained at a temperature of 36 ± 0.5 °C using a gravity-fed and heated perfusion system (custom built). A TILL Poly V light source (Till Photonics, Graefelfing, Germany) was used to excite Mitotracker (540 nm) and RNASelect^TM^ (475 nm) and 200 images at 1 Hz were recorded by a cooled CCD camera (PCO sensicam-QE) using TILL VisION (TILL Photonics) software. All image analysis was performed in Igor Pro (Wavemetrics, Portland, USA) using custom written routines (88) based on a previously described analysis algorithm (89). Neurites with at least one moving organelle were considered for the analysis. Spatio-temporal maps, or kymographs, were generated for each neuronal process in order to distinguish stationary mitochondria and moving mitochondria. The numbers of stationary and moving mitochondria were calculated by counting straight and diagonal lines on the kymographs.

In order to determine the directionality of the transport, cells were sparsely transduced with lentivirus expressing mitoDsRed2 at day 16 of the sMNs differentiation using an MOI of 0.5, optimized to visualize 1-2 labeled cells per field of view. Fluorescence live cell imaging of mitochondrial axonal transport was performed at day 38 of sMN differentiation at 63X magnification (Plan-Apochrat 1.40 NA oil DIC M27 objective, Carl Zeiss) using an Axio Observer Z1 inverted motorized microscope (Carl Zeiss) equipped with an Cy3 FL filter set (Carl Zeiss), Zen 2011 z-stack, time lapse and Definite focus modules (Carl Zeiss), and an S1 Environmental System incubation chamber (Carl Zeiss) maintained at 37 °C and 5% CO_2_. Media was changed 30 min prior to imaging to phenol-red Neurobasal supplemented with 1X GlutaMAX^TM^. Mitochondrial movements were recorded for 5 min using a 0.2 Hz capture of a 100 µm stretch of neurites and a small z-stack. At least 4 neurites were imaged per cell line per differentiation. sMN genotypes were counterbalanced and interleaved between experimental runs. Maximum intensity projections were computed in Fiji. Kymographs were generated and analyzed using KymoToolBox (90) in Fiji to determine the numbers of stationary (≤0.1 µm/s) versus motile mitochondria and the predominant directionality of movement (either away from (anterograde) or towards (retrograde) the soma).

### Proteins and dipeptide repeats

Full length human dynein with a SNAP_f_-tag on Dynein heavy chain was expressed in *Spodoptera frugiperda* Sf9 cells and purified and labelled with fluorophores as described previously (61, 91). SNAP_f_::BICD2N (not labelled with a fluorophore) was also produced from Sf9 cells using published methods (61). Native dynactin was purified from pig brain by SP-sepharose-based purification as described (92).

Active kinesin-1 (1-560 truncation containing the motor domain and dimerization stalk) was purified from Rosetta 2 *E. coli* (DE3, Novagen) transformed with the pET17-K560-GFP-His plasmid (59) and cultured at 37 °C. Expression was induced with 0.1 mM IPTG, with cells harvested 14-18 h later by centrifugation for 20 min at 3,000 x *g* (JLA 8.1 rotor) and resuspended in buffer A (50 mM Na-phosphate buffer (pH 8), 1 mM MgCl_2_, 250 mM NaCl, and 1 mM PMSF). Following lysis by sonication, insoluble material was removed by centrifugation for 35 min at 27,000 x *g* (70 Ti rotor). Initial purification was carried out by Ni^2+^-affinity chromatography (HisTrap^TM^FF 5 ml, GE Healthcare) and elution with 50 mM Na-phosphate buffer, 1 mM MgCl_2_, 250 mM NaCl, 250 mM imidazole, 50 mM Mg^2+^ ATP and 1 mM PMSF, pH 7.2. Protein was concentrated using Amicon 100 kDa 15 mL tubes at 4 °C (4000 x *g* spin) and washed in buffer A. Further purification was achieved through a microtubule pull down and release procedure, which enriches for the intact, active motor domain over degradation products (93). The protein sample was incubated with taxol-stabilized microtubules (15 mg mL^-1^) and 1 mM AMP-PNP (Sigma) and the mixture overlaid on a glycerol cushion (BRB80: 80 mM PIPES, pH 6.9, 1 mM EGTA, 1 mM MgCl_2_), 20 μM taxol and 60% glycerol (v/v)). Centrifugation was performed with a TLA 100 rotor at 317,000 x *g* for 10 min. The pellet was resuspended in BRB80 with 50 mM KCl and 5 mM ATP and incubated for 10 min to trigger release of microtubule-bound motors. Centrifugation at 317,000 x *g* for 10 min yielded supernatant containing the released active motors, which were snap frozen in small aliquots in 20% glycerol (v/v) and stored at −80 °C.

Porcine tubulin conjugated to fluorophores or biotin was purchased from Cytoskeleton, Inc. HPLC-purified dipeptide repeats were purchased from Pepscan and were either labelled with a single Cy5 dye at the N-terminus during synthesis or were labelled non-site-specifically in-house using Alexa488-Fluor® Protein Labeling Kit (Thermo Fisher Scientific) using published methods (94).

### Single molecule motility assays

Biotinylated and fluorescently-labelled porcine microtubules stabilized with taxol and GMP-CPP, PEG-biotin passivated glass surfaces, and motility chambers were prepared as described previously (91, 95). For assays with TMR-dynein, microtubules were polymerized in the presence of HiLyte488-tubulin, with the exception of the experiments documented in Fig. 3E, **Fig. S10B, C** and Fig. 4E-G, in which HiLyte647-tubulin was used for conditions in the presence of Alexa488-PR_30_. TRITC-tubulin was used for kinesin-1::GFP assays. Stabilized microtubules were immobilized on the biotinylated glass surface in the motility chamber for 30 s and subsequently washed once with motility buffer (30 mM HEPES, 5 mM MgSO4 1 mM EGTA, 1 mM DTT). For this and subsequent steps the pH of motility buffer was adjusted to 7.0 for kinesin-1 assays (kinesin-1 motility buffer) and 7.3 for dynein assays (dynein motility buffer), as this was necessary for optimal movement of the respective motors in the motility assay.

Kinesin-1::GFP dimers (96 nM) were stored on ice before motility assays in kinesin-1 motility buffer. For dynein assays, 138 nM TMR-SNAP::dynein dimers, 585 nM dynactin and 1.9 µM BICD2N dimers (molar ratio 1:4:14) were mixed on ice for 5-10 min in dynein motility buffer. An independent assembly of dynein, dynactin and BICD2N was used for each dynein assay chamber. Kinesin-1::GFP solutions were diluted 1 in 20 into kinesin-1 motility buffer in the presence or absence of DPRs at the final concentrations indicated in the Results section. Dynein, dynactin, BICD2N assembly mixtures were diluted 133-fold (with the exception of the complexes in Fig. 3C and **Fig. S11B** which were diluted 800-fold) in dynein motility buffer in the presence and absence of DPRs. Both final motility buffers also contained 25 mM KCl, 1 mg ml^−1^ α-casein, 5 mM MgATP, and an oxygen scavenging system (1.25 μM glucose oxidase, 140 nM catalase, 71 mM 2-mercaptoethanol, 25 mM glucose) to minimize photobleaching (96). Diluted protein samples were promptly injected into imaging chambers.

Imaging was performed at room temperature (23 ± 1 °C) with a Nikon TIRF microscope system controlled with Micro-Manager (97) and equipped with a 100X oil objective (Nikon APO TIRF, 1.49 NA oil), as well as with Coherent Sapphire 488 nm (150 mW), Coherent Sapphire 561 nm (150 mW), and Coherent CUBE 641 nm (100 mW) lasers. Images were acquired with an iXon^EM^+ DU-897E EMCCD camera (Andor), which gave pixel dimensions of 105 x 105 nm. Different channels were imaged sequentially through automated switching of emission filters (GFP, Cy3, and Cy5 (Chroma Technology Corp)). For each chamber, at least two dual-colour acquisitions of 300 frames were made at the maximum achievable frame rate (∼2 frames s^−1^) and 100 ms exposure per frame. Three chambers were imaged for every experimental condition with the exception of 1 µM PR30 in Fig. 3D and **Fig. S10A**, where one chamber was imaged.

Kymographs were generated and manually analyzed using Fiji (98). Microtubule position was determined by the fluorescent tubulin signal or a projection of motor or DPR signal over the duration of the movie. From each chamber, typically 10 microtubules were selected for analysis across two movies (five microtubules per movie), with the exception of Fig. 3E and **Fig. S10B** and **C** where 15 microtubules were selected across three movies when assaying PR_30_ addition. Microtubules were chosen before visualizing the motile properties of associated motor complexes or pattern of DPR association. Preference was given to microtubules that were relatively long and well separated from other microtubules. Subsequent analysis of kymographs was carried out by the researcher blind to the identity of the samples, with the exception of the experiments documented in Fig. 3D, E and **Fig. S10**. Blinding was not possible in these cases because either the magnitude of the effects on motility made it clear from which conditions the kymographs were derived or the batch of kymographs for analysis contained a single condition.

Interactions of fluorescently labelled motors with microtubules were operationally considered as binding events if they were ≥ 2 s (4 frames) in duration. As described previously (61), some processive complexes changed velocity within a run, which necessitated the calculation of velocities of individual constant-velocity segments. Run lengths were calculated from the total displacement of continuous movements of complexes, irrespective of changes in velocity. Pauses within or at the end of runs were scored only if the stationary event exceeded 1.5 s (≥ 3 frames). For Fig. 3C, background signal was quantified by generating kymographs from randomly selected microtubule-free regions of the coverslip of lengths equal to the median microtubule length of those used for analysis. Although kymographs displayed in Fig. 3A had background subtracted in Fiji with a rolling ball radius of 100 pixels for illustrative purposes, the raw data was used for all quantitative analysis.

### Quantification of *in vitro* microtubule binding by peptides

To quantify microtubule binding by Cy5-labelled DPRs, five TRITC-labelled microtubules were selected randomly from three images acquired from three independent chamber preparations. Mean Cy5 fluorescent intensity was then calculated using Fiji. Background signal was first subtracted with a rolling ball radius of 50 pixels to generate uniform fluorescent intensity across the region. Enrichment of fluorescence on microtubules was calculated by subtracting mean ‘background intensity’ from microtubule-associated intensity. Background intensity was calculated from five lines per image equivalent to the median length of microtubules assessed, which were positioned in microtubule-free areas close to the selected microtubules. Binding in Fig. 4A, B was quantified in the presence of the kinesin-1 complex and kinesin motility buffer; however, we verified in separate analyses that the presence of motors is not required for MT binding.

### Analysis of motor pausing and detachment on PR_30_-decorated microtubules

Single-molecule motility data for each motor was acquired as described above. Additional analysis of the co-localization of pause and detachment sites for kinesin-1 and dynein with intense PR_30_ puncta was carried out in Fiji. Co-incidence was scored if there was at least a 2-pixel overlap between the PR_30_ signal and signal from the incoming motor complex. For the simulated control, randomized patterns of PR_30_ foci were generated by overlaying the PR_30_ binding pattern from other microtubules within the same field of view onto the kymographs of motor events on the original microtubule. This procedure led to simulated PR_30_ banding patterns of equivalent density. Original and simulated images were assessed blind to their identities to avoid unconscious bias.

### Mouse husbandry and procedures

All mouse husbandry and procedures were performed in accordance with institutional guidelines and approved by the Stanford Administrative Panel on Animal Care (APLAC). C57BL/6 mice (Jackson Laboratory) were used. For spinal cord dissection, mice were anesthetized with isoflurane and transcardially perfused with 0.9% saline. Isolated spinal cords were used immediately for sample preparation.

### PR_30_ protein precipitation

Mouse spinal cords were disintegrated in PBS buffer with Halt Protease inhibitor (Thermo Fisher Scientific) with a Dounce homogenizer. The cell suspension was subsequently lysed using a probe sonicator (Branson Sonifier 250, VWR Scientific) on ice. The lysate was cleared from the insoluble fraction by centrifugation for 15 min at 10,000 rpm (11,292 x *g*) at 4 °C (Eppendorf 5427 R). The supernatant was retrieved and the procedure repeated until no pellet was visible after centrifugation. The protein concentration was measured using the Micro BCA assay (Thermo Fisher Scientific). PR_30_ (Pepscan) was added to a final concentration of 50 mM to 0.5 mg of soluble lysate in a total volume of 400 µl and samples were incubated for 15 min. The volume of the samples was increased to 1.5 ml with PBS, before gently spinning down the PR droplets at 4,000 rpm (3220 x *g*) for 5 min (Eppendorf 5810 R). Eppendorf tubes with phase separated samples were dispensed in 50 ml Falcon tubes and spun down in a swinging-bucket centrifuge. Pellets were subsequently washed with 1 ml PBS and vortexed before spinning down again. Washing steps were repeated three times. The resulting pellets were processed for LC-MS/MS.

For crosslinked samples (see **Table S5**), after incubation with PR_30_, paraformaldehyde (Sigma-Aldrich) was added to a final concentration of 0.5% for 5 min. 500 μl of 2 M Glycine (Sigma-Aldrich) was added for 5 min to quench paraformaldehyde. Samples were subsequently treated as uncrosslinked samples. Samples for analysis by SDS-PAGE and silver staining were generated identically. Silver staining (Thermo Fisher Scientific) was performed according to the manufacturer’s instructions.

### Proteomics sample preparation and LC-MS/MS analysis

Pellets were redissolved in 8 M urea in PBS, and solubilized by sonication. The total sample volume was 1.5 ml. Proteins in each sample were reduced with 5 mM DTT (30 min at 55 °C) and then alkylated by addition of 10 mM iodoacetamide for 15 min at room temperature in the dark. Samples were further diluted to a final urea concentration of 2 M and proteins digested with trypsin (Promega) (1/100, w/w) overnight at 37 °C. Peptides were then purified on Omix C18 tips (Agilent), dried and re-dissolved in solvent A (25 ml 0.1% trifluoroacetic acid (TFA) in dH_2_O/acetonitrile (98:2, v/v)) of which 10 ml was injected for LC-MS/MS analysis on an Ultimate 3000 RSLCnano System (Dionex, Thermo Fisher Scientific) in line connected to a Q Exactive HF mass spectrometer with a Nanospray Flex Ion source (Thermo Fisher Scientific). Trapping was performed at 10 ml/min for 4 min in solvent A (on a reverse phase column produced in-house, 100 mm I.D. x 20 mm, 5 mm beads C18 Reprosil-Pur, Dr. Maisch) followed by loading the sample on a 40-cm column packed in the needle (produced in-house, 75 mm I.D. 3 400 mm, 1.9 mm beads C18 Reprosil-HD, Dr. Maisch). Peptides were eluted by an increase in solvent B (0.1% formic acid in dH_2_O/acetonitrile (2:8, v/v)) in linear gradients from 2% to 3% in 100 min, then from 30% to 56% in 40 min and finally from 56% to 99% in 5 min, all at a constant flow rate of 250 nl/min. The mass spectrometer was operated in data-dependent mode, automatically switching between MS and MS/MS acquisition for the 16 most abundant ion peaks per MS spectrum. Full-scan MS spectra (375-1500 m/z) were acquired at a resolution of 60,000 after accumulation to a target value of 3,000,000 with a maximum fill time of 60 ms. The 16 most intense ions above a threshold value of 22,000 were isolated (window of 1.5 Th) for fragmentation at a normalized collision energy of 32% after filling the trap at a target value of 100,000 for maximum 45 ms. The S-lens RF level was set at 55 and precursor ions with single and unassigned charge states were excluded.

### Protein identification and quantification

Data analysis was performed with MaxQuant (version 1.5.3.30) (99) using the Andromeda search engine with default search settings including a false discovery rate set at 1% on both the peptide and protein level. Spectra were searched against the human proteins in the Uniprot/Swiss-Prot database (database release version of April 2016 containing 20,103 human protein sequences, www.uniprot.org). The mass tolerance for precursor and fragment ions were set to 4.5 and 20 ppm, respectively, during the main search. Enzyme specificity was set as C-terminal to arginine and lysine, also allowing cleavage at proline bonds with a maximum of three missed cleavages. Variable modifications were set to oxidation of methionine residues and acetylation of protein N-termini. Only proteins with at least one unique or razor peptide were retained leading to the identification of 1811 human proteins. Proteins were quantified by the MaxLFQ algorithm integrated in the MaxQuant software (100). A minimum ratio count of two unique or razor peptides was required for quantification. Further data analysis was performed with the Perseus software (version 1.5.3.0) after loading the protein groups file from MaxQuant. Proteins only identified by site and reverse database hits were removed. Missing values were imputed from a normal distribution around the detection limit.

### Co-Immunoprecipitation

Motor neurons were treated with 10 µM of HA-labelled DPR peptides (PR_20_, GR_20_, GP_20_; Pepscan) for 3 h. Subsequently, medium was removed and washed once with ice-cold DPBS (Sigma). Motor neurons were gently scraped off the dish and centrifuged for 5 min, 400 x *g* at 4 °C. Cell pellets were crosslinked using 1% PFA (Sigma) in DPBS (Sigma) and incubated for 10 min at room temperature. The crosslinking reaction was stopped with 1/10 volume of 1.25 M glycine pH 8 (Thermo Fisher Scientific). Crosslinked cell pellets were washed 3 times with ice-cold DPBS followed by lysis or snap freezing and storage at −80°C. Cell pellets were lysed in ice-cold lysis buffer (150 mM NaCl, 25 mM Tris, 1 mM EDTA, 1% NP40, 5% glycerol, pH 7.4) supplemented with complete EDTA-free protease inhibitor cocktail (Roche Diagnostics). Cell debris was removed via centrifugation and supernatants mixed with Pierce Anti-HA magnetic beads (Thermo Fisher Scientific). The mixture was incubated for 1 h at room temperature with agitation. Afterwards, the beads were washed 3 times with lysis buffer. The HA-tagged dipeptides and associated material were eluted using the manufacturer’s buffers.

For western blot analysis, samples were resolved on 4-20% mini-protean TGX stain-free gels (Biorad) and transferred to nitrocellulose membrane (Biorad). Membrane was blocked with 5% non-fat milk (Biorad) in Tris Buffer Saline Solution with 0.1% tween (TBST) for 2 h at room temperature and incubated with primary antibodies overnight at 4 °C (**Table S7**). The next day the membranes were washed 3 times with TBST and incubated for 1 h with secondary Trueblot antibodies (Rockland). Proteins were detected by enhanced chemiluminescence reagents (Thermo Fisher Scientific).

### Proximity ligation assay

Duolink® proximity ligation assay (PLA®) (Sigma-Aldrich, DUO92101) was performed according to the manufacturer’s instructions. Briefly, samples were fixed with 4% paraformaldehyde in PBS for 20 min, rinsed three times and incubated for 1 h at room temperature in PBS containing 0.2% Triton X-100 and 5% donkey serum. Cells were incubated overnight at 4 °C in 2% donkey serum containing the primary antibodies. After washing twice in 1x Duolink wash buffer A for 5 min, the cells were incubated with secondary antibodies conjugated with oligonucleotides (PLA probes anti-mouse MINUS and anti-rabbit PLUS) for 1 h in a pre-heated humidity chamber at 37 °C. See **Table S7** for details of antibodies. The samples were washed twice in Duolink washing buffer A for 5 min and then the Duolink ligation solution was applied to the samples for 30 min in a pre-heated humidity chamber at 37 °C. Cells were rinsed twice with Duolink washing buffer A for 5 min and subsequently the Duolink amplification-polymerase solution was applied to the samples in a dark pre-heated humidity chamber for 100 min at 37 °C. The samples were then washed twice in 1x Duolink washing buffer B for 10 min at room temperature followed by a 1 min wash with 0.01x Duolink washing buffer B. The cells were stained with Phalloidin for 1 h in PBS with 2% donkey serum, rinsed three times with PBS and incubated for 20 min in PBS/DAPI (NucBlue™ Live ReadyProbes™ Reagent, Invitrogen, R37605). Coverslips were mounted using ProLong™ Gold antifade reagent (Invitrogen, P36934) and confocal images obtained using a Leica SP8 DMI8 confocal microscope. PLA signals were recognized as red fluorescent spots.

### Immunofluorescence of post-mortem brains

To assess co-localization between GR aggregates and KIF5B in *C9orf72* post-mortem brains we made use of the central nervous system (CNS) tissues previously collected in the UZ Leuven brain biobank in accordance with the ethics review board upon written informed consent. In this study, two *C9orf72* expansion carriers (one ALS case and one ALS/FTD case) and two controls (ALS patients without the *C9orf72* expansion) were used (**Table S1**). Frozen frontal cortex slides of 7-8 µm thickness were washed with the BondTM Wash Solution (Leica Biosystems, AR9590) for 15 min at room temperature and stained overnight with the primary antibodies at 4 °C (Kinesin heavy chain: Millipore MAB1614 1:50; Poly-GR: Millipore MABN778 1:50). The following day, slides were washed three times with the BondTM Wash Solution before the secondary antibodies was added for 90 min at room temperature. After three washing steps with the BondTM Wash Solution, the slides were mounted using ProLong™ Gold antifade reagent with DAPI (Invitrogen, P36935). Confocal images were obtained using a Leica SP8 DMI8 confocal microscope. Images were analyzed, formatted and quantified with ImageJ software.

### *Drosophila* husbandry and strains

Flies were fed with Iberian food (5.5% (w/v) glucose, 5% (w/v) baker’s yeast, 3.5% (w/v) organic flour, 0.75% (w/v) agar, 16.4 mM Methyl-4-hydroxybenzoate, 0.004% (v/v) propionic acid) and maintained at 25 °C on a 12-h-light, 12-h-dark cycle at 50±5% humidity. The following strains were used: *yw* (control (Bullock lab stocks)); *GMR-Gal4 UAS-GR_36_/CyO* and *GMR-Gal4 UAS-PR_36_/CyO* (20); *P[Khc+]* (*Khc* genomic rescue construct pk9a (64)); FRT *G13 Khc^27^/CyO* (*Khc^null^ allele*) (101).

### Assessment of *Drosophila* viability and eye defects

Male *GMR-Gal4 UAS-DPR* (*GR_36_* or *PR_36_*) flies were mated with females with altered *Khc* gene dosage or *yw* controls. Although *GMR-Gal4* is predominantly expressed in the eye, its ability to cause lethality in combination with *UAS-GR_36_* (20) indicates that it has some activity in other tissues (the eye is not required for survival to adulthood). Each type of cross was set up in triplicate, with at least two egg-lays (of three days each) analyzed per cross. Adult offspring were collected and aged to 4-6 days post-eclosion before analysis. For all crosses that produced a sufficient number of offspring (> 15 internal control animals that did not inherit the *GMR-Gal4 UAS-DPR* chromosome) we calculated the fraction of male flies of different genotypes that inherited the *GMR-Gal4 UAS-DPR* chromosome (male flies were used for all phenotypic analysis to control for any sex-specific phenotypic effects). These values were plotted as a percentage of the expected fraction for each genotype assuming no lethality.

For analysis of eye defects, a single eye per DPR-expressing male was scored using a qualitative scale of eye disruption (no, mild, moderate, severe, very severe, or total pigment loss; **Table S6**). Necrotic patches were not classed as pigmentation. See Fig. 6B and **Fig. S15A** for example images for each category. All eye scoring was carried out blind of the genotypes to avoid unconscious bias. In control experiments, we confirmed that altering *Khc* gene dosage does not affect survival or eye morphology in the absence of expression of *GR_36_* or *PR_36_* (**Table S6**).

### *Drosophila* eye imaging

Example images of the different classes of eye phenotype were acquired using a Nikon ’Multiphot’ macro photography set up, with a 6X Macro-Nikkor 35 mm lens and a Sony a6300 ’mirrorless’ 24 megapixel camera, followed by processing in Adobe Photoshop CS6.

### Statistics

Statistical analyses and data plotting were performed using Prism 8 (GraphPad). Normality of the data was test using D’Agostino & Pearson normality test. Data containing more than two groups showing normal distribution were analyzed with a one-way ANOVA test and Tukey post hoc test. Data which were not normally distributed were analyzed using the Kruskal-Wallis non-parametric test with Dunn’s correction for multiple comparisons. Data with two groups showing normal distribution were analyzed using Student’s t-test; otherwise, the Mann–Whitney U test was used. * p<0.05, ** p<0.01, *** p<0.001, **** p<0.0001 were considered significant. Data values represent mean ± SD, unless indicated otherwise. For the *in vitro* motility assays a two-tailed Welch’s t-test was used when comparing two groups with Gaussian data distribution with unequal variance. A one-way ANOVA test with Dunnett’s correction was used for multiple comparisons to the control or Tukey’s correction when comparing between all conditions. Fisher’s Exact test was performed when comparing proportions of summed categorical variables.

## ACKNOWLEDGEMENTS

This work was supported by grants from KU Leuven (C1-C14-17-107), Opening the Future Fund (KU Leuven), the Alzheimer Research Foundation (SAO-FRA n° 2017/023), the Flemish Government initiated Flanders Impulse Program on Networks for Dementia Research (VIND n°135043), the Fund for Scientific Research Flanders (FWO Vlaanderen n° G.0909.15, G0F8516N, G0B2819N), the Agency for Innovation by Science and Technology (IWT n° 150031, 135043), the European E-Rare-3 project INTEGRALS, the ALS Liga België, the National Lottery of Belgium and the KU Leuven funds “Een Hart voor ALS”, “Laeversfonds voor ALS Onderzoek” and “Valéry Perrier Race against ALS Fund”. P.V.D. holds a senior clinical investigatorship of FWO Vlaanderen and is supported through the E. von Behring Chair for Neuromuscular Disorders. F.L.Y., V.M. and S.L.B. are supported by the UK Medical Research Council (file reference number MC_U105178790). S.B. is supported by an EMBO Long-Term Fellowship. A.R.M. is a Lady Edith Wolfson Clinical Fellow and is jointly funded by the Medical Research Council and the Motor Neurone Disease Association (MR/R001162/1). The Chandran Lab receives funding from the UK Dementia Research Institute partner funders: the UK Medical Research Council, Alzheimer’s Research UK, and the Alzheimer’s Society. L.D. is funded by a PhD Fellowship of the Research Foundation–Flanders (FWO-Vlaanderen) (1165119N). We would like to acknowledge members of the Bullock and the Neurobiology lab for support and advice throughout. In addition we would like to thank: Andrew Carter and Ahmet Yildiz labs for reagents and advice on the purification of recombinant kinesin-1, Werend Boesmans for advice on transport measurement, Francis Impens from the VIB proteomics expertise center for help with the mass spec analysis, Joke Terryn for the help with the Operetta CLS High Content Screening (HCS) System, Rosanna K. Ma for the technical assistance with the mice and Don Mahad for providing the mito-DsRed2 plasmid.

## AUTHOR CONTRIBUTIONS/COMPETING INTERESTS

L.F. and F.L.Y. designed and performed most experiments, analyzed the data and wrote the paper. S.B. performed the PR_30_ spinal cord interactome experiment and wrote the paper. M.D.D. helped perform DPR pulldowns. A.M., T.S.B. and S.C. provided the *C9orf72* and corrected isogenic cell lines, and A.M. performed transport experiments with them. A.S. performed and analyzed patch-clamp data. R.F., W.G., J.B. and T.V. contributed to iPSC culture. M.M. helped with the statistical analysis. L.D., D.R.T. and K.P. provided the post-mortem material. L.D. and D.R.T. helped perform, and provided inputs on, post-mortem experiments. M.v.B. performed the Southern blot experiment. D.R. and A.M. performed the ELISA experiment. A.D.G. supervised the PR_30_ spinal cord interactome experiment. P.K. provided iPSC lines. P.V.B. assisted with the intracellular transport experiments. V.M., C.V. and L.V.D.B. provided critical feedback and inputs into the project. S.L.B. and P.V.D. supervised the project, designed experiments, discussed the data and wrote the paper. All authors approved the final manuscript. The authors declare no competing interests.

## REFERENCES

1. A. M. DeJesus-Hernandez, I. R. Mackenzie, B. F. Boeve, A. L. Boxer, M. Baker, N. J. Rutherford, A. M. Nicholson, N. C. A. Finch, H. Flynn, J. Adamson, N. Kouri, A. Wojtas, P. Sengdy, G. Y. R. Hsiung, A. Karydas, W. W. Seeley, K. A. Josephs, G. Coppola et al., Expanded GGGGCC hexanucleotide repeat in noncoding region of *C9ORF72* causes chromosome 9p-Linked FTD and ALS. Neuron 72, 245–256 (2011).

2. B. A. E. Renton, E. Majounie, A. Waite, J. Simon-Sanchez, S. Rollinson, J. R. Gibbs, J. C. Schymick, H. Laaksovirta, J. C. van Swieten, L. Myllykangas, H. Kalimo, A. Paetau, Y. Abramzon, A. M. Remes, A. Kaganovich, S. W. Scholz, J. Duckworth, J. Ding et al., A hexanucleotide repeat expansion in *C9ORF72* is the cause of chromosome 9p21-linked ALS-FTD. Neuron 72, 257–268 (2011).

3. A. J. Waite, D. Bäumer, S. East, J. Neal, H. R. Morris, O. Ansorge, and D. J. Blake, Reduced C9orf72 protein levels in frontal cortex of amyotrophic lateral sclerosis and frontotemporal degeneration brain with the *C9ORF72* hexanucleotide repeat expansion. Neurobiol. Aging 35, 1779.e5–1779.e13 (2014).

4. J. Cooper-Knock, M. J. Walsh, A. Higginbottom, J. R. Highley, M. J. Dickman, D. Edbauer, P. G. Ince, S. B. Wharton, S. A. Wilson, J. Kirby, G. M. Hautbergue, and P. J. Shaw, Sequestration of multiple RNA recognition motif-containing proteins by *C9orf72* repeat expansions. Brain 137, 2040–2051 (2014).

5. C. J. Donnelly, P. W. Zhang, J. T. Pham, A. R. Heusler, N. A. Mistry, S. Vidensky, E. L. Daley, E. M. Poth, B. Hoover, D. M. Fines, N. Maragakis, P. J. Tienari, L. Petrucelli, B. J. Traynor, J. Wang, F. Rigo, C. F. Bennett, S. Blackshaw et al., RNA toxicity from the ALS/FTD *C9ORF72* expansion is mitigated by antisense intervention. Neuron 80, 415–428 (2013).

6. A. R. Haeusler, C. J. Donnelly, G. Periz, E. A. J. Simko, P. G. Shaw, M. S. Kim, N. J. Maragakis, J. C. Troncoso, A. Pandey, R. Sattler, J. D. Rothstein, and J. Wang, *C9orf72* nucleotide repeat structures initiate molecular cascades of disease. Nature 507, 195–200 (2014).

7. Y. B. Lee, H. J. Chen, J. N. Peres, J. Gomez-Deza, J. Attig, M. Štalekar, C. Troakes, A. L. Nishimura, E. L. Scotter, C. Vance, Y. Adachi, V. Sardone, J. W. Miller, B. N. Smith, J. M. Gallo, J. Ule, F. Hirth, B. Rogelj et al., Hexanucleotide repeats in ALS/FTD form length-dependent RNA foci, sequester RNA binding proteins, and are neurotoxic. Cell Rep. 5, 1178–1186 (2013).

8. K. Mori, S. Lammich, I. R. A. Mackenzie, I. Forné, S. Zilow, H. Kretzschmar, D. Edbauer, J. Janssens, G. Kleinberger, M. Cruts, J. Herms, M. Neumann, C. Van Broeckhoven, T. Arzberger, and C. Haass, HnRNP A3 binds to GGGGCC repeats and is a constituent of p62-positive/TDP43-negative inclusions in the hippocampus of patients with *C9orf72* mutations. Acta Neuropathol. 125, 413–423 (2013).

9. Z. Xu, M. Poidevin, X. Li, Y. Li, L. Shu, D. L. Nelson, H. Li, C. M. Hales, M. Gearing, T. S. Wingo, and P. Jin, Expanded GGGGCC repeat RNA associated with amyotrophic lateral sclerosis and frontotemporal dementia causes neurodegeneration. Proc. Natl. Acad. Sci. 110, 7778–7783 (2013).

10. P. E. A. Ash, K. F. Bieniek, T. F. Gendron, T. Caulfield, W. Lin, M. Dejesus-hernandez, M. M. Van Blitterswijk, K. Jansen-west, J. W. P. Iii, R. Rademakers, K. B. Boylan, D. W. Dickson, and L. Petrucelli, Unconventional translation of *C9ORF72* GGGGCC expansion generates insoluble polypeptides specific to c9FTD / ALS. Neuron 77, 639–646 (2013).

11. K. Mori, S. M. Weng, T. Arzberger, S. May, K. Rentzsch, E. Kremmer, B. Schmid, H. A. Kretzschmar, M. Cruts, C. Van Broeckhoven, C. Haass, and D. Edbauer, The *C9orf72* GGGGCC repeat is translated into aggregating dipeptide-repeat proteins in FTLD/ALS. Science (80-.). 339, 1335–1338 (2013).

12. K. Mori, T. Arzberger, F. A. Grässer, I. Gijselinck, S. May, K. Rentzsch, S. M. Weng, M. H. Schludi, J. Van Der Zee, M. Cruts, C. Van Broeckhoven, E. Kremmer, H. A. Kretzschmar, C. Haass, and D. Edbauer, Bidirectional transcripts of the expanded *C9orf72* hexanucleotide repeat are translated into aggregating dipeptide repeat proteins. Acta Neuropathol. 126, 881–893 (2013).

13. T. Zu, Y. Liu, M. Bañez-coronel, T. Reid, O. Pletnikova, and J. Lewis, RAN proteins and RNA foci from antisense transcripts in *C9ORF72* ALS and frontotemporal dementia. Proc. Natl. Acad. Sci. 110, 4968–4977 (2013).

14. X. Wen, W. Tan, T. Westergard, K. Krishnamurthy, S. S. Markandaiah, Y. Shi, S. Lin, N. A. Shneider, J. Monaghan, U. B. Pandey, P. Pasinelli, J. K. Ichida, and D. Trotti, Antisense proline-arginine RAN dipeptides linked to C9ORF72-ALS/FTD form toxic nuclear aggregates that initiate in vitro and in vivo neuronal death. Neuron 84, 1213–1225 (2014).

15. S. Almeida, E. Gascon, H. Tran, H. J. Chou, T. F. Gendron, S. Degroot, A. R. Tapper, C. Sellier, N. Charlet-Berguerand, A. Karydas, W. W. Seeley, A. L. Boxer, L. Petrucelli, B. L. Miller, and F. B. Gao, Modeling key pathological features of frontotemporal dementia with *C9ORF72* repeat expansion in iPSC-derived human neurons. Acta Neuropathol. 126, 385–399 (2013).

16. D. Sareen, J. G. O’Rourke, P. Meera, A. K. M. G. Muhammad, S. Grant, M. Simpkinson, S. Bell, S. Carmona, L. Ornelas, A. Sahabian, T. Gendron, L. Petrucelli, M. Baughn, J. Ravits, M. B. Harms, F. Rigo, C. F. Bennett, T. S. Otis et al., Targeting RNA foci in iPSC-derived motor neurons from ALS patients with a *C9ORF72* repeat expansion. Sci. Transl. Med. 5, 1–13 (2013).

17. S. B. Yamada, T. F. Gendron, T. Niccoli, N. R. Genuth, R. Grosely, Y. Shi, I. Glaria, N. J. Kramer, L. Nakayama, S. Fang, T. J. I. Dinger, A. Thoeng, G. Rocha, M. Barna, J. D. Puglisi, L. Partridge, J. K. Ichida, A. M. Isaacs et al., RPS25 is required for efficient RAN translation of *C9orf72* and other neurodegenerative disease-associated nucleotide repeats. Nat. Neurosci. 22, 1383–1388 (2019).

18. R. Lopez-gonzalez, D. Yang, M. Pribadi, T. S. Kim, G. Krishnan, and S. Yoen, Partial inhibition of the overactivated Ku80-dependent DNA repair pathway rescues neurodegeneration in *C9ORF72*-ALS / FTD. Proc. Natl. Acad. Sci. 116, 9628–9633 (2019).

19. I. Kwon, S. Xiang, M. Kato, L. Wu, P. Theodoropoulos, T. Wang, J. Kim, J. Yun, Y. Xie, and S. L. McKnight, Poly-dipeptides encoded by the *C9orf72* repeats bind nucleoli, impede RNA biogenesis, and kill cells. Science (80-.). 345, 1139–1146 (2014).

20. S. Mizielinska, S. Grönke, T. Niccoli, C. E. Ridler, E. L. Clayton, A. Devoy, T. Moens, F. E. Norona, I. O. C. Woollacott, J. Pietrzyk, K. Cleverley, A. J. Nicoll, S. Pickering-brown, J. Dols, M. Cabecinha, O. Hendrich, P. Fratta, E. M. C. Fisher et al., *C9orf72* repeat expansions cause neurodegeneration in *Drosophila* through arginine-rich proteins. Science (80-.). 345, 1192–1195 (2014).

21. B. D. Freibaum, Y. Lu, R. Lopez-Gonzalez, N. C. Kim, S. Almeida, K.-H. Lee, N. Badders, M. Valentine, B. L. Miller, P. C. Wong, L. Petrucelli, H. J. Kim, F.-B. Gao, and J. P. Taylor, GGGGCC repeat expansion in *C9orf72* compromises nucleocytoplasmic transport. Nature 525, 129–133 (2015).

22. A. Jovičič, J. Mertens, S. Boeynaems, E. Bogaert, N. Chai, S. B. Yamada, J. W. Paul, S. Sun, J. R. Herdy, G. Bieri, N. J. Kramer, F. H. Gage, L. Van Den Bosch, W. Robberecht, and A. D. Gitler, Modifiers of *C9orf72* dipeptide repeat toxicity connect nucleocytoplasmic transport defects to FTD/ALS. Nat. Neurosci. 18, 1226–1229 (2015).

23. L. H. Tran, S. Almeida, J. Moore, T. F. Gendron, U. D. Chalasani, Y. Lu, X. Du, J. A. Nickerson, L. Petrucelli, Z. Weng, and F. B. Gao, Differential toxicity of nuclear RNA foci versus dipeptide repeat proteins in a Drosophila model of C9ORF72 FTD/ALS. Neuron 87, 1207–1214 (2015).

24. D. Yang, A. Abdallah, Z. Li, Y. Lu, S. Almeida, and F. B. Gao, FTD/ALS-associated poly(GR) protein impairs the Notch pathway and is recruited by poly(GA) into cytoplasmic inclusions. Acta Neuropathol. 130, 525–535 (2015).

25. M. S. Boeynaems, E. Bogaert, E. Michiels, I. Gijselinck, A. Sieben, A. Jovičić, G. De Baets, W. Scheveneels, J. Steyaert, I. Cuijt, K. J. Verstrepen, P. Callaerts, F. Rousseau, J. Schymkowitz, M. Cruts, C. Van Broeckhoven, P. Van Damme, A. D. Gitler et al., *Drosophila* screen connects nuclear transport genes to DPR pathology in c9ALS/FTD. Sci. Rep. 6, 7–14 (2016).

26. N. J. Kramer, M. S. Haney, D. W. Morgens, A. Jovičić, J. Couthouis, A. Li, J. Ousey, R. Ma, G. Bieri, C. K. Tsui, Y. Shi, N. T. Hertz, M. Tessier-lavigne, J. K. Ichida, M. C. Bassik, and A. D. Gitler, CRISPR–Cas9 screens in human cells and primary neurons identify modifiers of C9ORF72 dipeptide-repeat-protein toxicity. Nat. Genet. 50, 603–612 (2018).

27. B. Swinnen, A. Bento-Abreu, T. F. Gendron, S. Boeynaems, E. Bogaert, R. Nuyts, M. Timmers, W. Scheveneels, N. Hersmus, J. Wang, S. Mizielinska, A. M. Isaacs, L. Petrucelli, R. Lemmens, P. Van Damme, L. Van Den Bosch, and W. Robberecht, A zebrafish model for *C9orf72* ALS reveals RNA toxicity as a pathogenic mechanism. Acta Neuropathol. 135, 427–443 (2018).

28. R. Lopez-Gonzalez, Y. Lu, T. F. Gendron, A. Karydas, H. Tran, D. Yang, L. Petrucelli, B. L. Miller, S. Almeida, and F. B. Gao, Poly(GR) in *C9ORF72*-related ALS/FTD compromises mitochondrial function and increases oxidative stress and DNA damage in iPSC-derived motor neurons. Neuron 92, 383–391 (2016).

29. S. Y. Choi, R. Lopez-gonzalez, G. Krishnan, H. L. Phillips, A. N. Li, W. W. Seeley, W. Yao, S. Almeida, and F. Gao, *C9ORF72*-ALS/FTD-associated poly(GR) binds Atp5a1 and compromises mitochondrial function *in vivo*. Nat. Neurosci. 22, 851–862 (2019).

30. S. Yin, R. Lopez-Gonzalez, R. C. Kunz, J. Gangopadhyay, C. Borufka, S. P. Gygi, F. B. Gao, and R. Reed, Evidence that *C9ORF72* dipeptide repeat proteins associate with U2 snRNP to cause mis-splicing in ALS/FTD patients. Cell Rep. 19, 2244–2256 (2017).

31. K. Kanekura, T. Yagi, A. J. Cammack, J. Mahadevan, M. Kuroda, M. B. Harms, T. M. Miller, and F. Urano, Poly-dipeptides encoded by the *C9ORF72* repeats block global protein translation. Hum. Mol. Genet. 25, 1803–1813 (2016).

32. H. Hartmann, D. Hornburg, M. Czuppa, J. Bader, M. Michaelsen, D. Farny, T. Arzberger, M. Mann, F. Meissner, and D. Edbauer, Proteomics and *C9orf72* neuropathology identify ribosomes as poly-GR/PR interactors driving toxicity. Life Sci. Alliance 1, 1–13 (2018).

33. T. G. Moens, T. Niccoli, K. M. Wilson, M. L. Atilano, and N. Birsa, *C9orf72* arginine-rich dipeptide proteins interact with ribosomal proteins in vivo to induce a toxic translational arrest that is rescued by eIF1A. Acta Neuropathol. 137, 487–500 (2019).

34. V. Lafarga, O. Sirozh, I. Díaz-López, M. Hisaoka, E. Zarzuela, J. Boskovic, B. Jovanovic, R. Fernandez-Leiro, J. Muñoz, G. Stoecklin, I. Ventoso, and O. Fernandez-Capetillo, Generalized displacement of DNA- and RNA-binding factors mediates the toxicity of arginine-rich cell-penetrating peptides. Biorxiv, doi.org/10.1101/441808 (2018).

35. K. Zhang, C. J. Donnelly, A. R. Haeusler, J. C. Grima, J. B. Machamer, P. Steinwald, E. L. Daley, S. J. Miller, K. M. Cunningham, S. Vidensky, S. Gupta, M. A. Thomas, I. Hong, S. L. Chiu, R. L. Huganir, L. W. Ostrow, M. J. Matunis, J. Wang et al., The *C9orf72* repeat expansion disrupts nucleocytoplasmic transport. Nature 525, 56–61 (2015).

36. S. Boeynaems, E. Bogaert, D. Kovacs, and P. Van Damme, Phase separation of *C9orf72* dipeptide repeats perturbs stress granule dynamics. Mol. Cell 65, 1044–1055 (2017).

37. K. H. Lee, P. Zhang, H. J. Kim, D. M. Mitrea, M. Sarkar, B. D. Freibaum, J. Cika, M. Coughlin, J. Messing, A. Molliex, B. A. Maxwell, N. C. Kim, J. Temirov, J. Moore, R. M. Kolaitis, T. I. Shaw, B. Bai, J. Peng et al., *C9orf72* dipeptide repeats impair the assembly, dynamics, and function of membrane-less organelles. Cell 167, 774–788 (2016).

38. Y. Lin, E. Mori, M. Kato, S. Xiang, L. Wu, I. Kwon, and S. L. McKnight, Toxic PR poly-dipeptides encoded by the *C9orf72* repeat expansion target LC domain polymers. Cell 167, 789–802 (2016).

39. A. E. Renton, A. Chiò, and B. J. Traynor, State of play in amyotrophic lateral sclerosis genetics. Nat. Neurosci. 17, 17–23 (2014).

40. B. N. Smith, N. Ticozzi, C. Fallini, A. S. Gkazi, S. Topp, K. P. Kenna, E. L. Scotter, J. Kost, P. Keagle, J. W. Miller, D. Calini, C. Vance, E. W. Danielson, C. Troakes, C. Tiloca, S. Al-Sarraj, E. A. Lewis, A. King et al., Exome-wide rare variant analysis identifies *TUBA4A* mutations associated with familial ALS. Neuron 84, 324–331 (2014).

41. A. Nicolas, K. P. Kenna, A. E. Renton, N. Ticozzi, F. Faghri, R. Chia, J. A. Dominov, B. J. Kenna, M. A. Nalls, P. Keagle, A. M. Rivera, W. van Rheenen, N. A. Murphy, J. J. F. A. van Vugt, J. T. Geiger, R. van der Spek, H. A. Pliner, Shankaracharya et al., Genome-wide analyses identify *KIF5A* as a novel ALS gene. Neuron 97, 1268–1283 (2018).

42. C. Münch, R. Sedlmeier, T. Meyer, V. Homberg, A. D. Sperfeld, A. Kurt, J. Prudlo, G. Peraus, C. O. Hanemann, G. Stumm, and A. C. Ludolph, Point mutations of the p150 subunit of *dynactin* (DCTN1) gene in ALS. Neurology 63, 724–727 (2004).

43. S. Millecamps and and J.-P. Julien, Axonal transport deficits and neurodegenerative diseases. Nat. Rev. Neurosci. 14, 161–176 (2013).

44. K. J. De Vos and M. Hafezparast, Neurobiology of axonal transport defects in motor neuron diseases: Opportunities for translational research? Neurobiol. Dis. 105, 283–299 (2017).

45. W. Guo, M. Naujock, L. Fumagalli, T. Vandoorne, P. Baatsen, R. Boon, L. Ordovás, A. Patel, M. Welters, T. Vanwelden, N. Geens, T. Tricot, V. Benoy, J. Steyaert, C. Lefebvre-Omar, W. Boesmans, M. Jarpe, J. Sterneckert et al., HDAC6 inhibition reverses axonal transport defects in motor neurons derived from *FUS*-ALS patients. Nat. Commun. 8, 1–14 (2017).

46. K. R. Baldwin, V. K. Godena, V. L. Hewitt, and A. J. Whitworth, Axonal transport defects are a common phenotype in *Drosophila* models of ALS. Hum. Mol. Genet. 25, 2378–2392 (2016).

47. M. Naumann, A. Pal, A. Goswami, X. Lojewski, J. Japtok, A. Vehlow, M. Naujock, R. Günther, M. Jin, N. Stanslowsky, P. Reinhardt, J. Sterneckert, M. Frickenhaus, F. Pan-Montojo, E. Storkebaum, I. Poser, A. Freischmidt, J. H. Weishaupt et al., Impaired DNA damage response signaling by FUS-NLS mutations leads to neurodegeneration and FUS aggregate formation. Nat. Commun. 9, 1–17 (2018).

48. N. Kreiter, A. Pal, X. Lojewski, P. Corcia, M. Naujock, P. Reinhardt, J. Sterneckert, S. Petri, F. Wegner, A. Storch, and A. Hermann, Age-dependent neurodegeneration and organelle transport deficiencies in mutant TDP43 patient-derived neurons are independent of TDP43 aggregation. Neurobiol. Dis. 115, 167–181 (2018).

49. B. T. Selvaraj, M. R. Livesey, C. Zhao, J. M. Gregory, O. T. James, E. M. Cleary, A. K. Chouhan, A. B. Gane, E. M. Perkins, O. Dando, S. G. Lillico, Y. B. Lee, A. L. Nishimura, U. Poreci, S. Thankamony, M. Pray, N. A. Vasistha, D. Magnani et al., C9ORF72 repeat expansion causes vulnerability of motor neurons to Ca^2+^-permeable AMPA receptor-mediated excitotoxicity. Nat. Commun. 9, 1–14 (2018).

50. T. G. Moens, S. Mizielinska, T. Niccoli, J. S. Mitchell, A. Thoeng, C. E. Ridler, S. Grönke, J. Esser, A. Heslegrave, H. Zetterberg, L. Partridge, and A. M. Isaacs, Sense and antisense RNA are not toxic in *Drosophila* models of *C9orf72*-associated ALS/FTD. Acta Neuropathol. 135, 445–457 (2018).

51. N. H. Alami, R. B. Smith, M. A. Carrasco, L. A. Williams, C. S. Winborn, S. S. W. Han, E. Kiskinis, B. Winborn, B. D. Freibaum, A. Kanagaraj, A. J. Clare, N. M. Badders, B. Bilican, E. Chaum, S. Chandran, C. E. Shaw, K. C. Eggan, T. Maniatis et al., Axonal transport of TDP-43 mRNA granules is impaired by ALS-causing mutations. Neuron 81, 536–543 (2014).

52. J. P. Taylor, R. H. Brown, and D. W. Cleveland, Decoding ALS: from genes to mechanism. Nature 539, 197–206 (2016).

53. A. D. Pilling, D. Horiuchi, C. M. Lively, and W. M. Saxton, Kinesin-1 and dynein are the primary motors for fast transport of mitochondria in *Drosophila* motor axons. Mol. Biol. Cell 17, 2057–2068 (2006).

54. M. Van Spronsen, M. Mikhaylova, J. Lipka, M. A. Schlager, D. J. Van Den Heuvel, M. Kuijpers, P. S. Wulf, N. Keijzer, J. Demmers, L. C. Kapitein, D. Jaarsma, H. C. Gerritsen, A. Akhmanova, and C. C. Hoogenraad, TRAK/Milton motor-adaptor proteins steer mitochondrial trafficking to axons and dendrites. Neuron 77, 485–502 (2013).

55. W. M. Saxton and P. J. Hollenbeck, The axonal transport of mitochondria. J. Cell Sci. 125, 2095–2104 (2012).

56. B. Ma, J. N. Savas, M. S. Yu, B. P. Culver, M. V. Chao, and N. Tanese, Huntingtin mediates dendritic transport of *β-actin* mRNA in rat neurons. Sci. Rep. 1, 1–11 (2011).

57. Y. Kanai, N. Dohmae, and N. Hirokawa, Kinesin transports RNA: isolation and characterization of an RNA-transporting granule. Neuron 43, 513–525 (2004).

58. P. K. Sahoo, D. S. Smith, N. Perrone-Bizzozero, and J. L. Twiss, Axonal mRNA transport and translation at a glance. J. Cell Sci. 131, 1–8 (2018).

59. G. Woehlke, A. K. Ruby, C. L. Hart, B. Ly, N. Hom-Booher, and R. D. Vale, Microtubule interaction site of the kinesin motor. Cell 90, 207–216 (1997).

60. R. J. Mckenney, W. Huynh, M. E. Tanenbaum, G. Bhabha, and R. D. Vale, Activation of cytoplasmic dynein motility by dynactin-cargo adapter complexes. Science (80-.). 345, 337–342 (2014).

61. M. A. Schlager, H. T. Hoang, L. Urnavicius, S. L. Bullock, and A. P. Carter, *In vitro* reconstitution of a highly processive recombinant human dynein complex. EMBO J. 33, 1855–1868 (2014).

62. S. Fredriksson, M. Gullberg, J. Jarvius, C. Olsson, K. Pietras, S. M. Gústafsdóttir, A. Östman, and U. Landegren, Protein detection using proximity-dependent DNA ligation assays. Nat. Biotechnol. 20, 473–477 (2002).

63. O. Soderberg, M. Gullberg, M. Jarvius, K. Ridderstrale, K.-J. Leuchowius, J. Jarvius, K. Wester, P. Hydbring, F. Bahram, L.-G. Larsson, and U. Landegren, Direct observation of individual endogenous protein complexes *in situ* by proximity ligation. Nat. Methods 3, 995–1000 (2006).

64. W. M. Saxton, J. Hicks, L. S. E. Goldstein, and C. Raff, Kinesin heavy chain is essential for viability and neuromuscular functions in *Drosophila*, but mutants show no defects in mitosis. Cell 64, 1093–1102 (1991).

65. A. Vagnoni and S. L. Bullock, A cAMP/PKA/Kinesin-1 axis promotes the axonal transport of mitochondria in aging Drosophila neurons. Curr. Biol. 28, 1265–1272 (2018).

66. A. Vagnoni, P. C. Hoffmann, and S. L. Bullock, Reducing Lissencephaly-1 levels augments mitochondrial transport and has a protective effect in adult Drosophila neurons. J. Cell Sci. 1, 178–190 (2016).

67. A. L. Jolly and V. I. Gelfand, Bidirectional intracellular transport: utility and mechanism. Biochem Soc Trans. 39, 1126–1130 (2012).

68. W. O. Hancock, Bidirectional cargo transport: moving beyond tug of war. Nat. Rev. Mol. Cell Biol. 15, 615–628 (2014).

69. K. Ikenaka, K. Kawai, M. Katsuno, Z. Huang, Y. Jiang, and Y. Iguchi, *dnc-1 dynactin 1* knockdown disrupts transport of autophagosomes and induces motor neuron degeneration. PLoS One 8, 1–18 (2013).

70. J. R. Levy, C. J. Sumner, J. P. Caviston, M. K. Tokito, S. Ranganathan, L. A. Ligon, K. E. Wallace, B. H. Lamonte, G. G. Harmison, I. Puls, K. H. Fischbeck, and E. L. F. Holzbaur, A motor neuron disease–associated mutation in p150Glued perturbs dynactin function and induces protein aggregation. J. Cell Biol. 172, 733–745 (2006).

71. J. K. Moore, D. Sept, and J. A. Cooper, Neurodegeneration mutations in dynactin impair dynein-dependent nuclear migration. Proc. Natl. Acad. Sci. 106, 5147–5152 (2009).

72. T. E. Lloyd, J. Machamer, K. O. Hara, J. H. Kim, S. E. Collins, M. Y. Wong, B. Sahin, W. Imlach, Y. Yang, E. S. Levitan, B. D. Mccabe, and A. L. Kolodkin, The p150^Glued^ CAP-Gly domain regulates initiation of retrograde transport at synaptic termini. Neuron 74, 344–360 (2012).

73. A. J. Moughamian and E. L. F. Holzbaur, Dynactin is required for transport initiation from the distal axon. Neuron 74, 331–343 (2012).

74. R. Balendra and A. M. Isaacs, C9orf72-mediated ALS and FTD: multiple pathways to disease. Nat. Rev. Neurol. 14, 544–558 (2018).

75. R. Dixit, J. L. Ross, Y. E. Goldman, and E. L. F. Holzbaur, Differential regulation of dynein and kinesin motor proteins by Tau. Science (80-.). 319, 1086–1090 (2008).

76. R. Tan, A. J. Lam, T. Tan, J. Han, D. W. Nowakowski, M. Vershinin, S. Simó, K. M. Ori-McKenney, and R. J. McKenney, Microtubules gate tau condensation to spatially regulate microtubule functions. Nat. Cell Biol. 21, 1078–1085 (2019).

77. M. Vershinin, B. C. Carter, D. S. Razafsky, S. J. King, and S. P. Gross, Multiple-motor based transport and its regulation by Tau. Proc. Natl. Acad. Sci. 104, 87–92 (2006).

78. A. R. Chaudhary, F. Berger, C. L. Berger, and A. G. Hendricks, Tau directs intracellular trafficking by regulating the forces exerted by kinesin and dynein teams. Traffic 19, 111–121 (2018).

79. V. Siahaan, J. Krattenmacher, A. A. Hyman, S. Diez, A. Hernández-vega, Z. Lansky, and M. Braun, Kinetically distinct phases of tau on microtubules regulate kinesin motors and severing enzymes. Nat. Cell Biol. 21, 1086–1092 (2019).

80. R. Rademakers, M. Cruts, and C. Van Broeckhoven, The role of Tau (MAPT) in frontotemporal dementia and related tauopathies. Hum. Mutat. 295, 277–295 (2004).

81. F. Paonessa, L. D. Evans, R. Solanki, J. Hardy, S. P. Jackson, F. J. Livesey, F. Paonessa, L. D. Evans, R. Solanki, D. Larrieu, S. Wray, and J. Hardy, Microtubules deform the nuclear membrane and disrupt nucleocytoplasmic transport in Tau-mediated frontotemporal dementia. Cell Rep. 26, 582–593 (2019).

82. J. A. Cross and M. P. Dodding, Motor–cargo adaptors at the organelle–cytoskeleton interface. Curr. Opin. Cell Biol. 59, 16–23 (2019).

83. S. L. Reck-Peterson, W. B. Redwine, R. D. Vale, and A. P. Carter, The cytoplasmic dynein transport machinery and its many cargoes. Nat. Rev. Mol. Cell Biol. 19, 382–398 (2018).

84. S. Saberi, J. E. Stauffer, J. Jiang, S. D. Garcia, E. Amy, D. Schulte, T. Ohkubo, C. L. Schloffman, M. Maldonado, M. Baughn, M. J. Rodriguez, D. Pizzo, D. Cleveland, and J. Ravits, Sense-encoded poly-GR dipeptide repeat proteins correlate to neurodegeneration and uniquely co-localize with TDP-43 in dendrites of repeat expanded *C9orf72* amyotrophic lateral sclerosis. Acta Neuropathol. 135, 459–474 (2018).

85. Y. Maury, R. A. Piskorowski, N. Salah-Mohellibi, V. Chevaleyre, M. Peschanski, C. Martinat, and P. Nedelec, Combinatorial analysis of developmental cues efficiently converts human pluripotent stem cells into multiple neuronal subtypes. Nat. Biotechnol. 33, 89–98 (2014).

86. C. Bock, E. Kiskinis, G. Verstappen, H. Gu, G. Boulting, Z. D. Smith, M. Ziller, G. F. Croft, M. W. Amoroso, D. H. Oakley, and A. Gnirke, Reference maps of human ES and iPS cell variation enable high-throughput characterization of pluripotent cell lines. Cell 144, 439–452 (2011).

87. M. Van Blitterswijk, M. Dejesus-hernandez, E. Niemantsverdriet, M. E. Murray, M. G. Heckman, N. N. Diehl, R. C. Petersen, B. F. Boeve, N. R. Graff, K. B. Boylan, L. Petrucelli, D. W. Dickson, and R. Rademakers, Association between repeat sizes and clinical and pathological characteristics in carriers of *C9ORF72* repeat expansions (Xpansize-72): a cross-sectional cohort study. Lancet Neurol. 12, 978–988 (2013).

88. C. D’Ydewalle, J. Krishnan, D. M. Chiheb, P. Van Damme, J. Irobi, A. P. Kozikowski, P. Vanden Berghe, V. Timmerman, W. Robberecht, and L. Van Den Bosch, HDAC6 inhibitors reverse axonal loss in a mouse model of mutant *HSPB1*–induced Charcot-Marie-Tooth disease. Nat. Med. 17, 968–974 (2011).

89. P. Vanden Berghe, G. W. Hennig, T. K. Smith, V. Berghe, G. W. Hennig, and K. Terence, Characteristics of intermittent mitochondrial transport in guinea pig enteric nerve fibers. Am J Physiol Gastrointest Liver Physiol 46, 671–682 (2004).

90. D. Zala, M. Hinckelmann, H. Yu, M. Menezes, S. Marco, and F. P. Cordelie, Vesicular glycolysis provides on-board energy for fast axonal transport. Cell 152, 479–491 (2013).

91. H. T. Hoang, M. A. Schlager, A. P. Carter, and S. L. Bullock, DYNC1H1 mutations associated with neurological diseases compromise processivity of dynein–dynactin–cargo adaptor complexes. Proc. Natl. Acad. Sci. 14, 1597–1606 (2017).

92. L. Urnavicius, K. Zhang, A. G. Diamant, C. Motz, M. A. Schlager, M. Yu, N. A. Patel, C. V. Robinson, and A. P. Carter, The structure of the dynactin complex and its interaction with dynein. Science (80-.). 347, 1441–1446 (2015).

93. M. Tomishige and R. D. Vale, Controlling kinesin by reversible disulfide cross-linking: identifying the motility-producing conformational change. J. Cell Biol. 151, 1081–1092 (2000).

94. S. Boeynaems, M. De Decker, P. Tompa, and L. Van Den Bosch, Arginine-rich peptides can actively mediate liquid-liquid phase separation. bio-protocol 7, 3–10 (2017).

95. M. A. Mcclintock, C. I. Dix, C. M. Johnson, S. H. Mclaughlin, R. J. Maizels, H. T. Hoang, and S. L. Bullock, RNA-directed activation of cytoplasmic dynein-1 in reconstituted transport RNPs. Elife 7, 1–29 (2018).

96. A. Yildiz, J. N. Forkey, S. A. Mckinney, T. Ha, Y. E. Goldman, and P. R. Selvin, Myosin V walks hand-over-hand: single fluorophore imaging with 1.5-nm localization. Science (80-.). 300, 2061–2066 (2003).

97. A. Edelstein, N. Amodaj, K. Hoover, R. Vale, and N. Stuurman, Computer control of microscopes using μManager. Curr. Protoc. Mol. Biol. 92, 1–18 (2010).

98. J. Schindelin, I. Arganda-carreras, E. Frise, V. Kaynig, M. Longair, T. Pietzsch, S. Preibisch, C. Rueden, S. Saalfeld, B. Schmid, J. Tinevez, D. J. White, V. Hartenstein, K. Eliceiri, P. Tomancak, and A. Cardona, Fiji: an open-source platform for biological-image analysis. Nat. Methods 9, 676–682 (2012).

99. J. Cox and M. Mann, MaxQuant enables high peptide identification rates, individualized p.p.b.-range mass accuracies and proteome-wide protein quantification. Nat. Biotechnol. 26, 1367–1372 (2008).

100. J. Cox, M. Y Hein, C. A. Luber, I. Paron, N. Nagaraj, and M. Mann, Accurate proteome-wide label-free quantification by delayed normalization and maximal peptide ratio extraction, termed MaxLFQ. Mol. Cell. Proteomics 13, 2513–2526 (2014).

101. K. M. Brendza, D. J. Rose, S. P. Gilbert, and W. M. Saxton, Lethal kinesin mutations reveal amino acids important for ATPase activation and structural coupling. J. Biol. Chem. 274, 31506–31514 (1999).

