## Supplementary Figures for "*C9orf72*-derived arginine-containing dipeptide repeats associate with axonal transport machinery and impede microtubule-based motility"

### SUPPLEMENTARY FIGURE LEGENDS

**Fig. S1: Generation and characterization of *C9orf72* and control iPSCs.** (A-C) Both control and *C9orf72* iPSCs expressed the pluripotency markers Nanog, Tra1-81, Oct4, Tra1-60, Sox2 and SSEA4. Scale bar, 100 μm. (D) hPSC ScoreCard results of trilineage differentiation potential confirmed that all generated iPSC lines have the capacity to differentiate towards all three germ layers (endoderm, mesoderm, ectoderm).

**Fig. S2: *C9orf72* repeat expansion analysis.** (A) Repeat length PCR shows that a repeat expansion is present in all the *C9orf72* iPSC lines and none of the controls from healthy individuals. All *C9orf72* lines are heterozygous for the repeat expansion. (B) Southern blot analysis of iPSCs and iPSC-derived sMNs estimating the size of the repeat expansion. C9-1B: 185, 540, 740 repeats; C9-2B: 770 repeats; C9-3B: 215 repeats. The presence of multiple repeat lengths in C9-1B might be due to instability of the expanded allele during iPSC reprogramming, as previously suggested (15).

**Fig. S3: Control and *C9orf72* iPSC lines differentiate into sMNs.** (A) Schematic overview of the motor neuron differentiation protocol. The experiments were performed at the end of the differentiation procedure (day 38), unless stated otherwise. (B, C) Both control and *C9orf72* lines efficiently differentiated into sMNs as demonstrated by the expression of ChAT, SMI32 (B) and ISLET1 (C) immunostaining. TUJ1 was used as a pan-neuronal marker. Scale bar, 50 μm. (D-E) Quantification of ChAT (D) and ISLET1 (E) positive cells expressed relative to the total number of DAPI labeled cells. Data are represented as mean ± S.D. Statistical significance was evaluated with an unpaired Student's t-test (ns, not significant; from N = 3 independent differentiations). Additional abbreviations: MNPs, motor neuron precursors; Y, Y-27632; SB, SB-431542; LDN, LDN-193189; CHIR, CHIR-99021; RA, retinoic acid; SAG, smoothened agonist; DAPT, N-[(3,5-Difluorophenyl)acetyl]-L-alanyl-2-phenylglycine-1,1-dimethylethyl ester; BDNF, brain-derived neurotrophic factor; GDNF, glial cell line-derived neurotrophic factor; CNTF, ciliary neurotrophic factor.

**Fig. S4: Electrophysiological characterization of iPSCs-derived sMNs.** (A) Representative trace of a mutant *C9orf72* (C9-2B) motor neuron exhibiting spontaneous action potentials recorded in current clamp mode without current injection. (B) Top panel: Action potentials are evoked by a series of current steps (0-60 pA, 10-pA increments, 1s duration). Lower panel: Representative traces of membrane potential responses to current steps recorded from a C9-2B motor neuron. The maximal number of action potentials in response to a depolarizing current step (red trace) was counted for every motor neuron. (C, D) Percentage of cells firing spontaneous (C) and evoked (D) action potentials (controls: n = 41; *C9orf72*: n = 52). (E, F) Firing frequency of spontaneous (E) and evoked (F) action potentials (spontaneous, control: n = 35; *C9orf72*: n = 41; evoked, control: n = 41; *C9orf72*: n = 51). Data are represented as mean ± S.D.; dots represent the number of patched cells. Statistical significance was evaluated with Mann-Whitney test (ns, not significant; from N = 3 independent differentiations).

**Fig. S5: Phenotypic characterization of the *C9orf72* lines.** (A-D) Digital Droplet PCR was used to calculate absolute levels of total *C9orf72* transcript variants (VALL) as well as its transcripts variants V1, V2, V3 in iPSCs and iPSC-derived sMNs. V1, V2 and V3 were normalized to total *C9orf72* levels; the total *C9orf72* levels were normalized to the *SCLY* reference gene. Data are represented as mean ± S.D. Statistical significance was evaluated with unpaired Student's t-test (\*\*\*\* $p < 0.0001$; \*$p < 0.05$; ns, not significant; from $N = 3$ independent experiments). (E) ELISA measurement of poly-GA and poly-GP dipeptides in controls and *C9orf72* derived sMNs. Data are represented as relative concentration with respect to a pool of healthy controls. Values represent mean of three independent measurements from $N = 3$ independent differentiations.

**Fig. S6: Additional data on the effect of the *C9orf72* hexanucleotide repeat expansion on mitochondria transport in sMNs.** (A) Examples of kymographs from each of the control and *C9orf72* patient cell lines. Y-axis, time; x-axis, distance; scale bars, 50 s and 30 $\mu\text{m}$. (B-D) Quantification of moving (B), stationary (C) and total (D) mitochondria normalized to 100 $\mu\text{m}$ neurite length. Total mitochondria are calculated as the sum of moving and stationary events. (E-G) Quantification of moving (E) stationary (F) and total (G) mitochondria normalized to 100 $\mu\text{m}$ neurite length at three different time points of the motor neuron differentiation. Data are represented as mean ± S.D.; dots represent the number of neurites measured (see **Table S3** for total numbers of neurites analyzed in each experiment). Statistical significance was evaluated with Mann–Whitney test (\*\*\*\* $p < 0.0001$; ns, not significant; from $N = 3$ independent differentiations).

**Fig. S7: Administration of synthetic DPRs to control MNs.** (A-C) DPRs were visualized by immunostaining in control sMNs after overnight treatment with two different doses of GP<sub>20</sub> (A), GR<sub>20</sub> (B) or PR<sub>20</sub> (C). Both GR<sub>20</sub> and PR<sub>20</sub>, due to their positive charge, were taken up by the cells. The uncharged peptide, GP<sub>20</sub>, was only able to pass the cellular membrane upon treatment of the cells with Streptolysin O (SLO). Cells treated with SLO without GP<sub>20</sub> were used as a control condition for this peptide, including for the transport analysis in **Fig. 2** and **Fig. S8**. For this specific experiment mCherry-GP<sub>20</sub> was used to visualize the peptide inside the cells while for all the other experiments a non-tagged GP<sub>20</sub> peptide was used. Scale bar, 10 $\mu\text{m}$. (D) Percentage of DRAQ7-positive cells after overnight treatment with increasing concentrations of GP<sub>20</sub>, GR<sub>20</sub> or PR<sub>20</sub>. Data are represented as mean ± S.E.M. Statistical significance was evaluated with one-way ANOVA multiple comparisons test with Tukey's correction (\*\*$p < 0.001$; \*$p < 0.01$; \*$p < 0.05$; ns, not significant; from $N = 2$ independent differentiations).

**Fig. S8: Additional data on the effect of the synthetic DPRs on transport of cargos in control sMNs.** (A-C) Quantification of moving (A) stationary (B) and total (C) mitochondria normalized to 100 $\mu\text{m}$ neurite length, upon treatment with GP<sub>20</sub>. (D-F) Quantification of moving (D), stationary (E) and total (F) mitochondria normalized to 100 $\mu\text{m}$ neurite length, upon treatment with GR<sub>20</sub>. (G-I) Quantification of moving (G), stationary (H) and total (I) mitochondria normalized to 100 $\mu\text{m}$ neurite length, upon treatment with PR<sub>20</sub>. (L-N) Quantification of moving (L), stationary (M) and total (N) RNA granules normalized to 100 $\mu\text{m}$ neurite length, upon treatment with PR<sub>20</sub>. Data are represented as mean ± S.D.; dots represent the number of neurites measured (see **Table S3** for total numbers of neurites analyzed in each experiment). Statistical significance was evaluated with Kruskal-Wallis test with Dunn's multiple comparison (\*\*\*\* $p < 0.0001$; \*\*\* $p < 0.001$; \*\* $p < 0.01$; ns, not significant; from $N = 3$ independent differentiations).

**Fig. S9: Additional data on the effect of PR<sub>30</sub> on mitochondria transport in control sMNs.** (A) Example kymographs of healthy MNs after treatment with PR<sub>30</sub>. Y-axis, time; x-axis, distance; scale bars, 50 s and 30 μm. (B-D) Quantification of moving (B), stationary (C) and total (D) mitochondria normalized to 100 μm neurite length, upon treatment with PR<sub>30</sub>. Data are represented as mean ± S.D.; dots represent the number of neurites measured (see **Table S3** for total numbers of neurites analyzed in each experiment). Statistical significance was evaluated with one-way ANOVA multiple comparisons test with Tukey's correction (G) (\*\*\* $p < 0.001$; \*\* $p < 0.01$; ns, not significant).

**Fig. S10: Quantification of the effects of varying PR chain length and concentration on additional motile properties of kinesin-1 and dynein.** (A, B) Quantification of effects of 1 μM or 10 μM PR<sub>30</sub> on additional motile properties of kinesin-1 (A) and dynein (B). (C, D) Quantification of effects of 10 μM PR<sub>20</sub> or PR<sub>30</sub> on motile properties of dynein (C) and kinesin-1 (D). In D, values for kinesin-1 in the presence of PR<sub>20</sub> and PR<sub>30</sub> are compared directly to their shared, internal control (no peptide), whereas in C data for PR<sub>20</sub> or PR<sub>30</sub> were acquired from two distinct experiments using dynein from independent protein purifications. Therefore, dynein's motile properties in the presence of PR<sub>20</sub> or PR<sub>30</sub> are compared to the corresponding internal control. In **Fig. 3E**, the control values for % processive complexes were pooled as they were very similar for the two experiments. Means ± S.D. are shown and dots represent values for individual microtubules. Statistical significance was evaluated with a one-way ANOVA multiple comparisons test with Tukey's correction (A, B, D) or an unpaired t test with Welch's correction (C) (\*\*\*\* $p < 0.0001$; \*\*\* $p < 0.001$; \*\* $p < 0.01$; \* $p < 0.05$; ns, not significant).

**Fig. S11: Quantification of the effect of 20-repeat dipeptides on additional motile properties of kinesin-1 and dynein.** (A, B) Quantification of effects of 10 μM GP<sub>20</sub>, GR<sub>20</sub> or PR<sub>20</sub> on the run length, run duration and total motor binding events for kinesin-1 (A) and dynein (B). Note that, unexpectedly, GP<sub>20</sub> has a statistically significant effect on the run duration and number of total binding events of dynein complexes. We also observe an increase in run length of kinesin-1 in the presence of GR<sub>20</sub> but not PR<sub>20</sub>. Means ± S.D. are shown and dots represent values for individual microtubules. Statistical significance was evaluated with a one-way ANOVA multiple comparisons test with Dunnett's correction (\*\*\*\* $p < 0.0001$; \*\*\* $p < 0.001$; \*\* $p < 0.01$; ns, not significant).

**Fig. S12: The punctate binding pattern of PR<sub>30</sub> on microtubules *in vitro* is observed at a range of peptide concentrations.** Example images of microtubules incubated with 1 and 10 μM PR<sub>30</sub> illustrating the presence of intense puncta in addition to continuous binding along the microtubule. Control and 10 μM PR<sub>30</sub> images are reproduced from **Fig 4A**. Arrowheads show examples of intense PR<sub>30</sub> puncta. Scale bar, 2.5 μm.

**Fig. S13: Additional data on interactors of arginine-rich DPRs.** (A, B) RNA transport granule components (A) and constituents of axonal transport cargos (B) were identified in the PR<sub>30</sub> interactome described in **Fig. 5A-D**. (C-E) Uncropped membranes from western blots related to **Fig. 5E**.

**Fig. S14: Supplementary figure 14: PR<sub>20</sub> localizes with β-III-Tubulin and KIF5B in sMNs.** (A) Immunostaining for PR<sub>20</sub> and β-III-Tubulin (TUJ1) within axonal growth cones of young sMNs (day 16) without and with treatment with synthetic PR<sub>20</sub> peptides. Arrowheads show co-localization of signals associated with microtubules. Scale bar, 5 μm. (B) Representative images showing positive proximity ligation assay (PLA) signals (present as red dots) for PR<sub>20</sub> and KIF5B in control sMNs after treatment with PR<sub>20</sub> synthetic peptides both in the cell body (upper panel) and along the neurites (lower panel). Scale bar, 15 μm.

**Fig. S15: Additional data on the ability kinesin-1 and dynein gene dosage to modify eye defects caused by poly-GR or poly-PR expression in *Drosophila*.** (A) Effect of Kinesin-1 heavy chain (Khc) gene dosage on the eye defects caused by GR<sub>36</sub> overexpression. Left, representative images of phenotypic categories. Scale bar, 100 μm. Right, quantification of eye phenotypes (total number of eyes analyzed is shown above bars) for GR<sub>36</sub> alone and GR<sub>36</sub> with overexpression of Khc (*P[Khc<sup>+</sup>]*) from a transgene containing the genomic locus. Due to the high incidence of lethality when GR<sub>36</sub> was combined with *Khc<sup>null</sup>/+* (**Fig. 6A**), there were insufficient progeny for meaningful analysis of eye phenotypes. Left chart shows overall data. Middle and right charts illustrate statistical evaluations of the proportion of eyes exhibiting the mildest (middle) and most severe (right) phenotypes upon Khc overexpression. (B) Statistical evaluations for effects of Khc gene dosage on all categories of eye defect severity caused by PR<sub>36</sub> overexpression. See **Fig. 6B** for collated data. In A and B, statistical significance was evaluated using Fisher's exact test. (\*\*\*\**p*<0.0001; \*\*\**p*<0.001; ns, not significant). (C) Pooled data from **Fig. 6B** broken down into individual egg lays. Note the high degree of consistency observed between equivalent egg lays. Asterisks for egg lays 2 and 5 of *P[Khc<sup>+</sup>]* highlight a relatively low number of offspring, which means that the values obtained are less likely to be representative of the general population. All flies were analyzed blind to the genotype.

### SUPPLEMENTARY MATERIALS

Supplementary tables (in Excel format) can be downloaded here:

https://www2.mrc-lmb.cam.ac.uk/groups/sbullock/Fumagalli_Young_Boeynaems_et_al_Tables/

**Table S1:** Clinical information of *C9orf72* iPSC lines and human autopsy cases.

**Table S2:** Passive membrane properties of control and *C9orf72* iPSC-derived sMNs.

**Table S3:** Total numbers of neurites analyzed in each experiment.

**Table S4:** Total numbers of motor complexes or microtubules analyzed in each experiment.

**Table S5:** List of identified proteins, GO analysis, and disease association from PR<sub>30</sub> interactome studies (see 'contents' tab for detailed information).

**Table S6:** Analysis of *Drosophila* control genotypes and descriptions of qualitative eye scoring categories depicted in **Fig. 6B** and **Fig. S15**.

**Table S7:** List of antibodies and stains used in this study.

Fig.S1

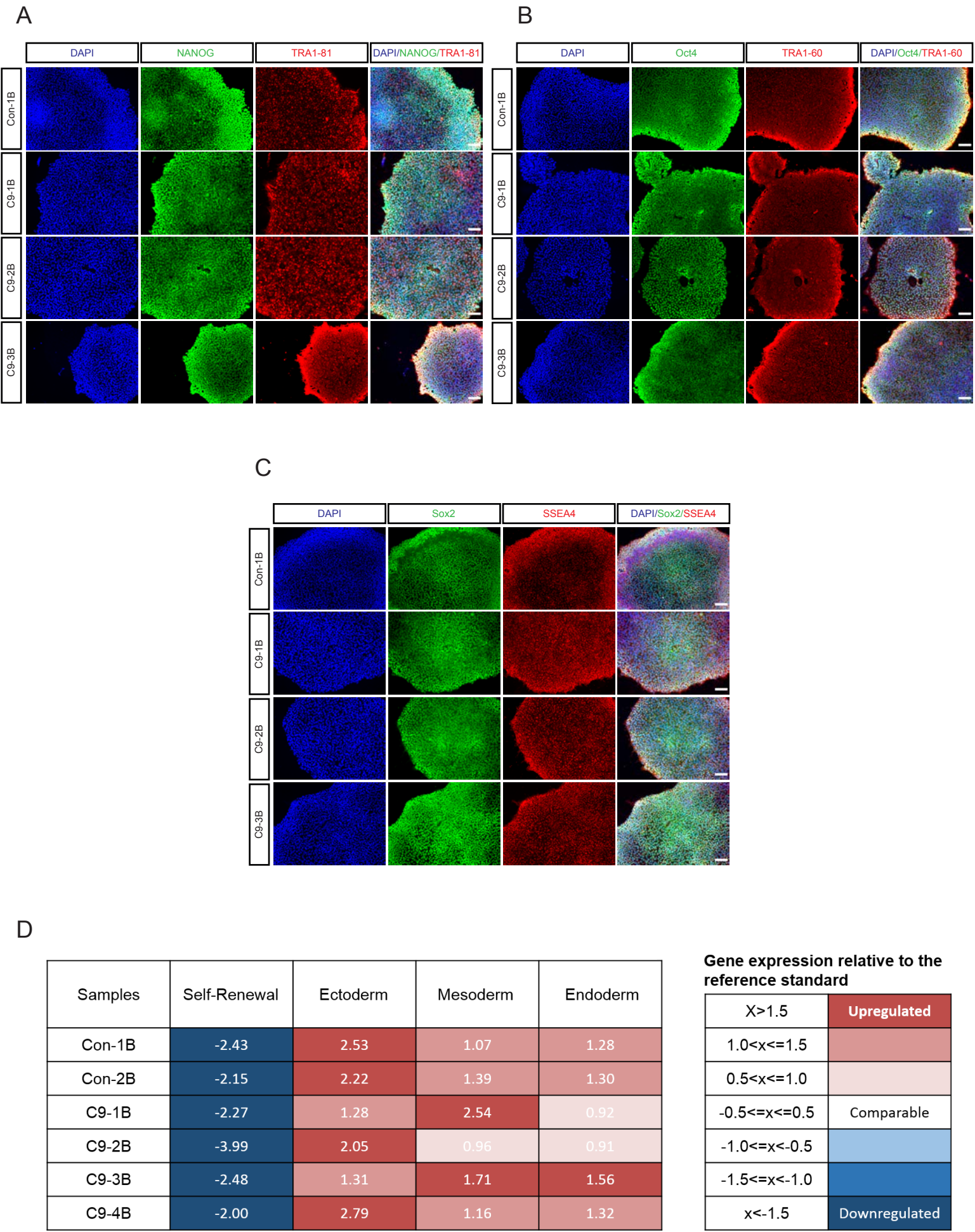

Fig.S2

A

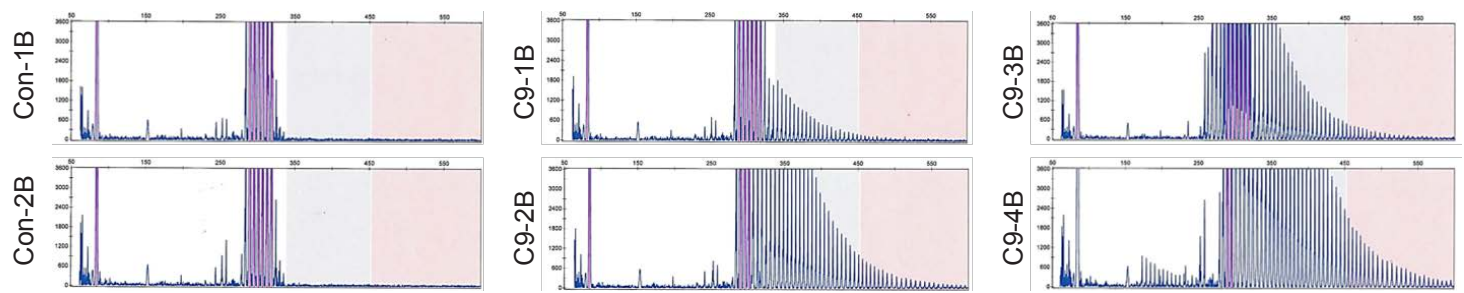

B

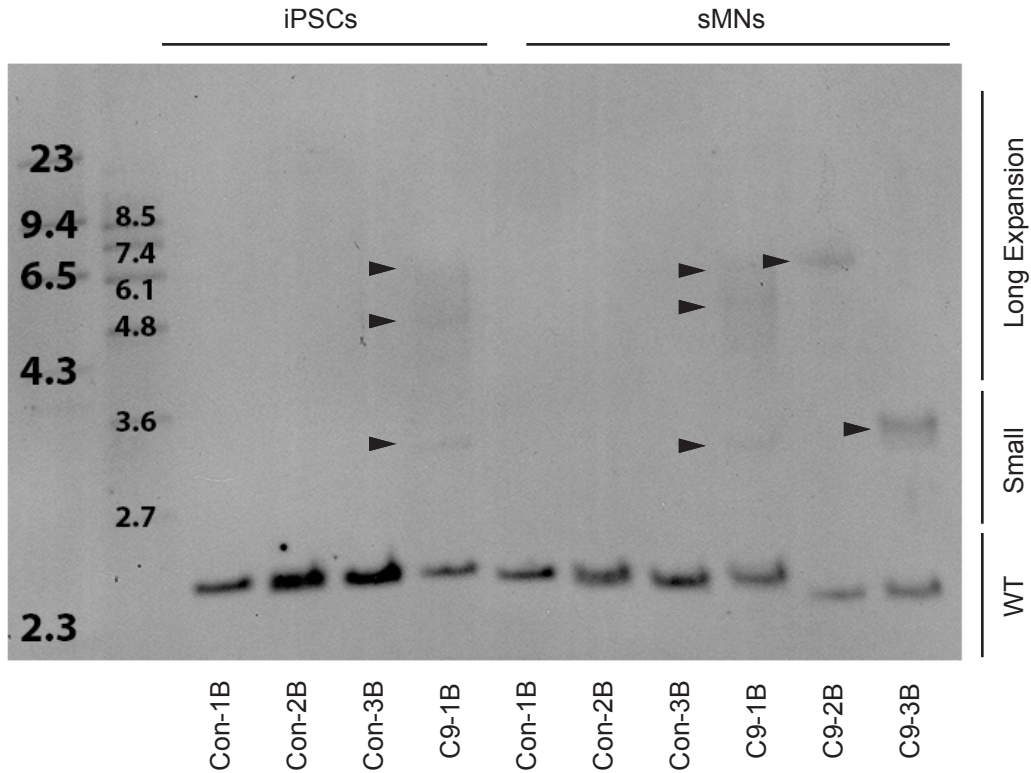

Fig.S3

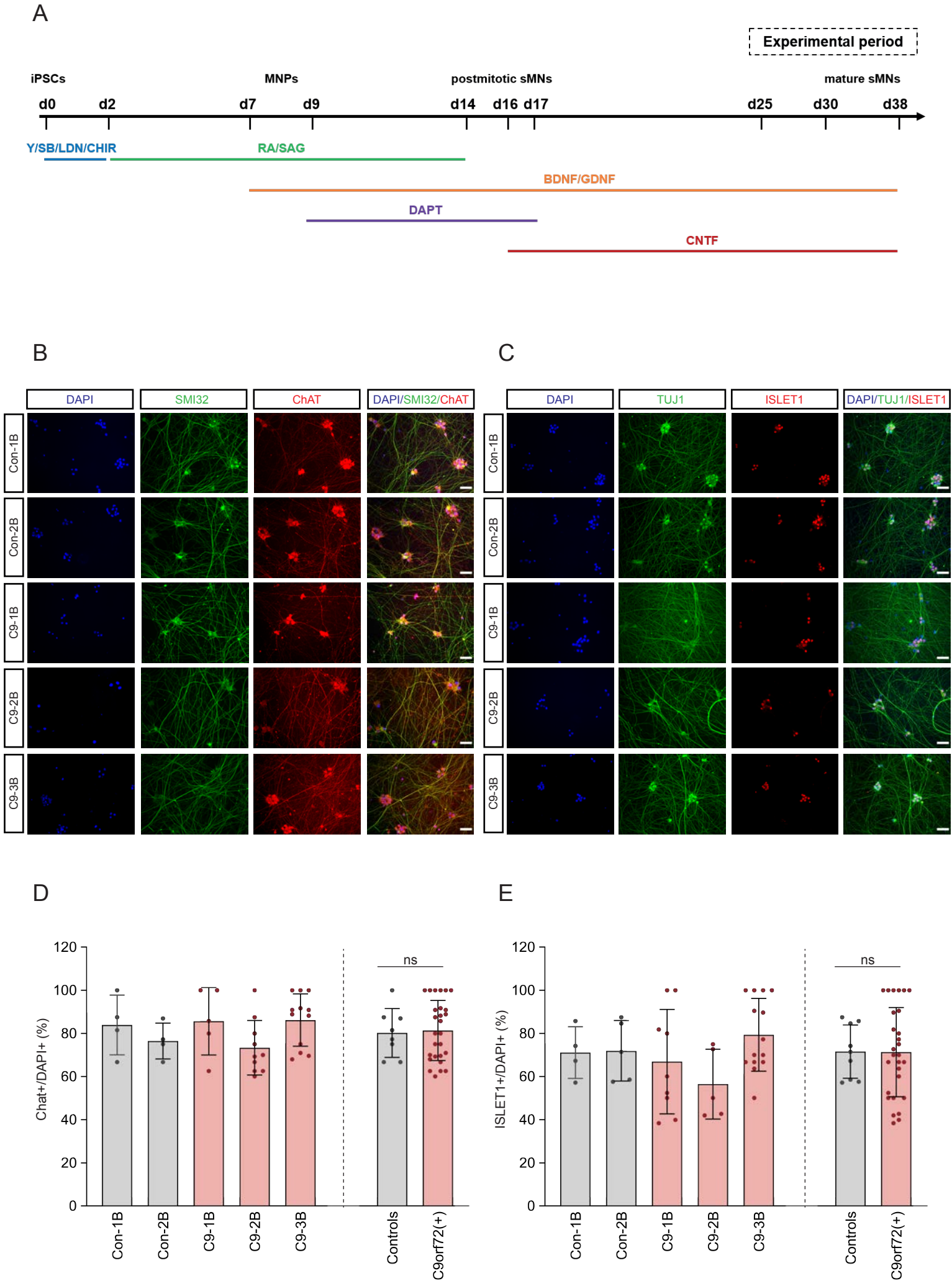

Fig.S4

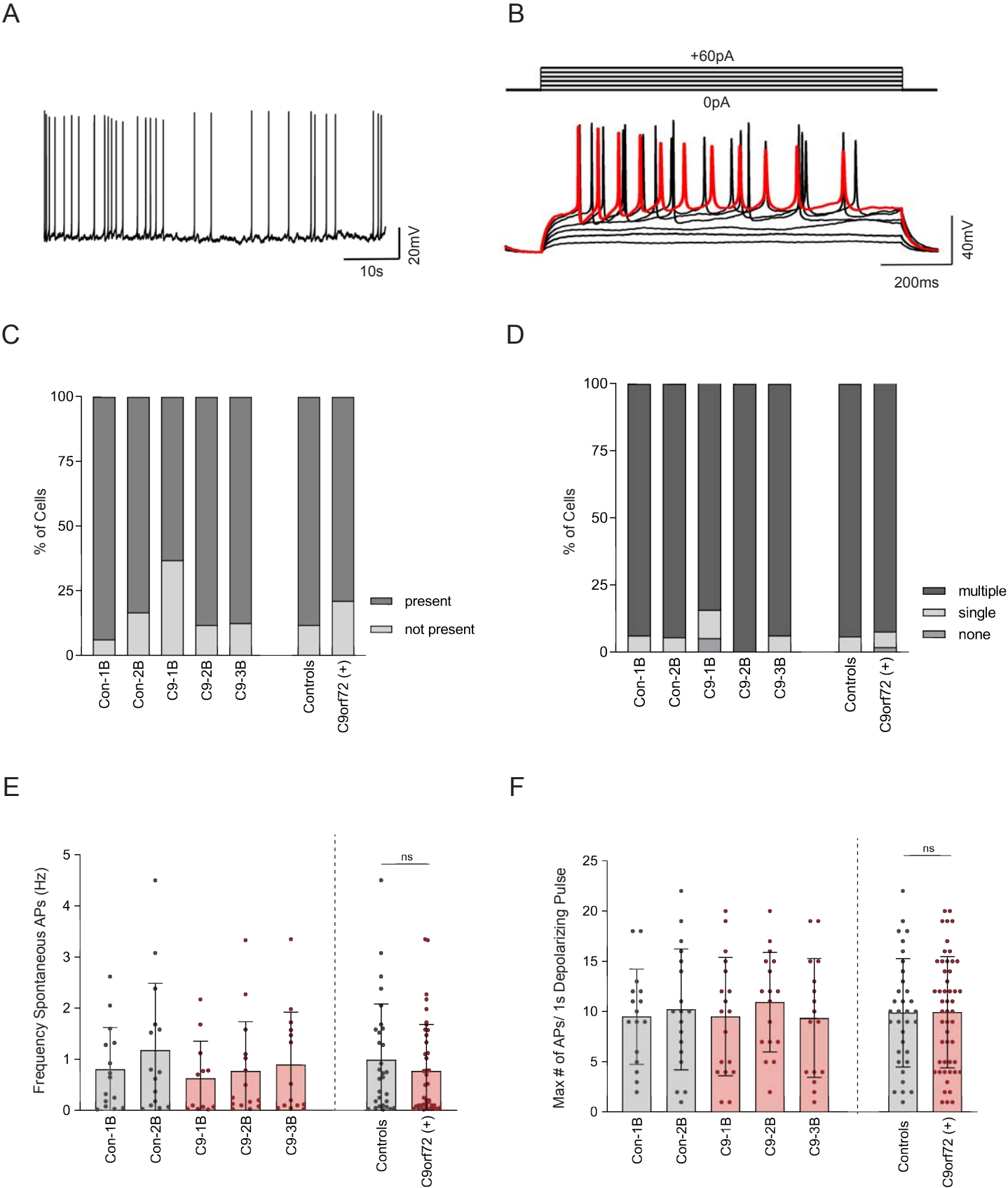

Fig.S5

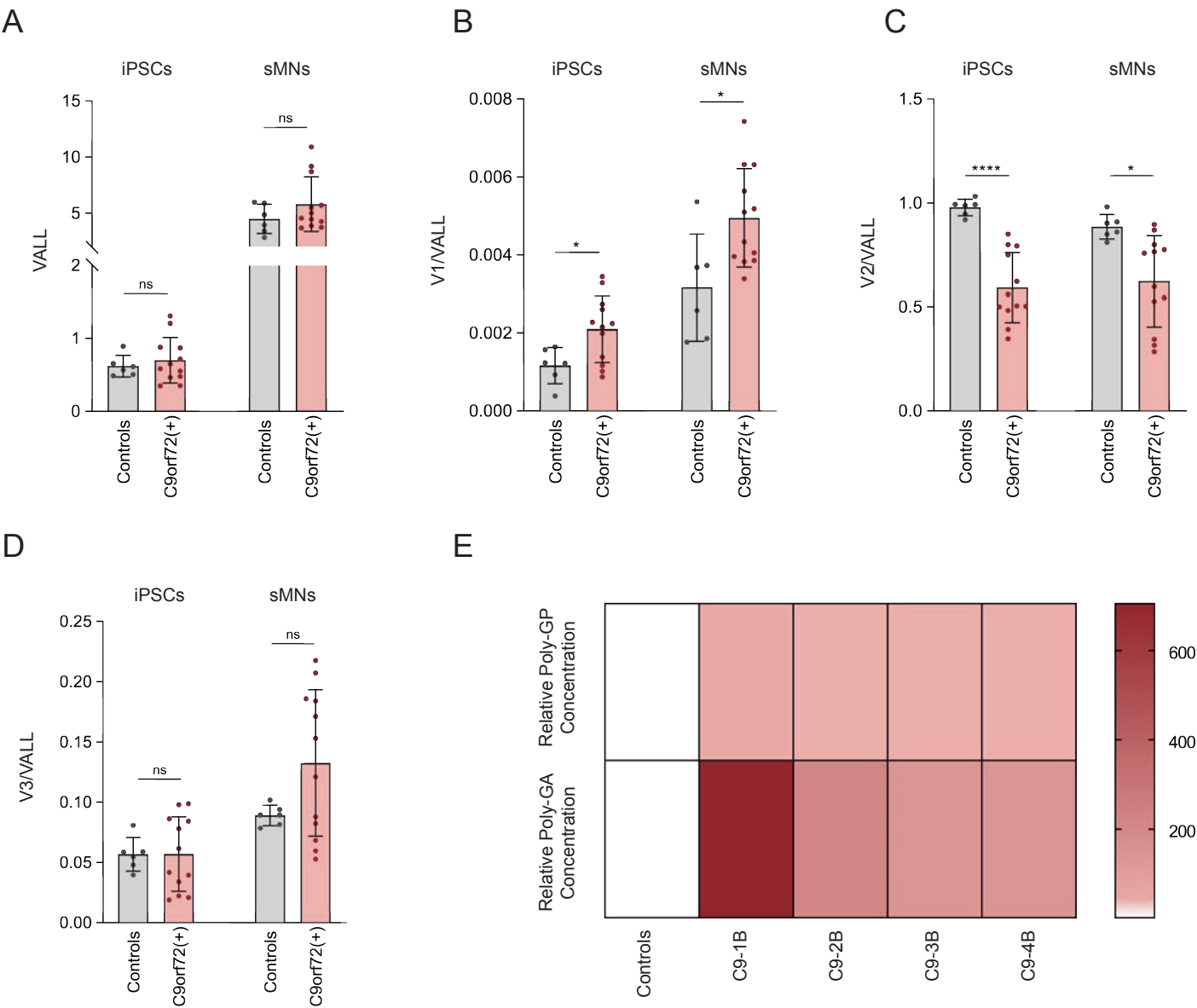

Fig.S6

A

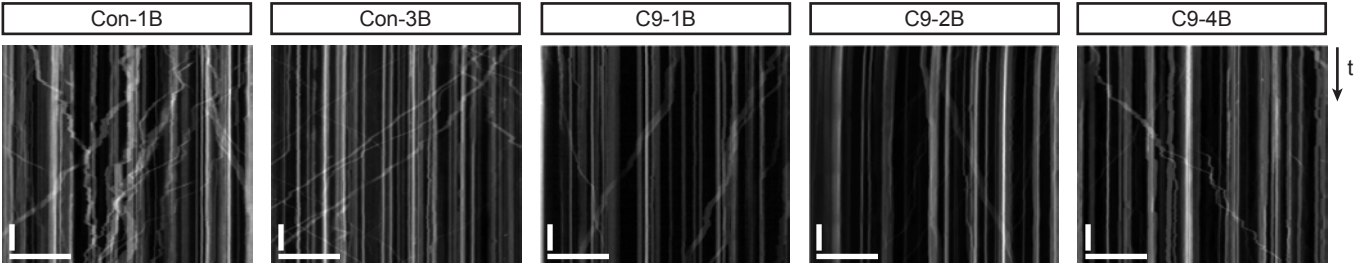

B

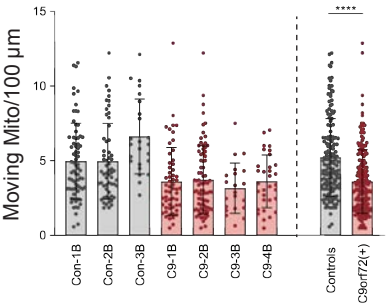

C

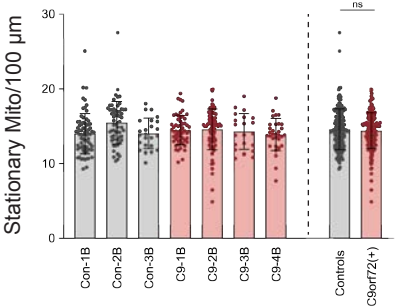

D

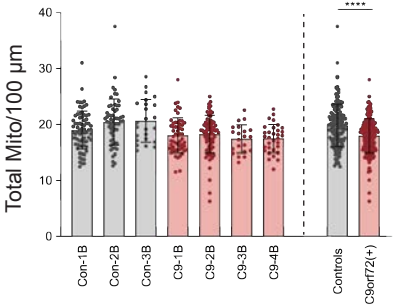

E

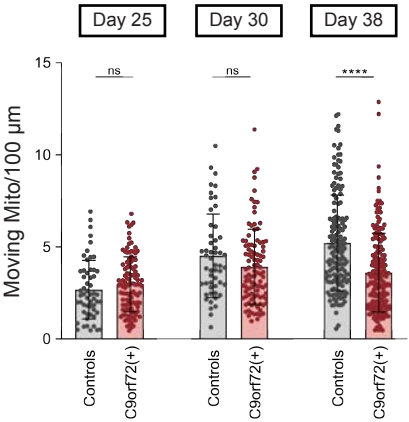

F

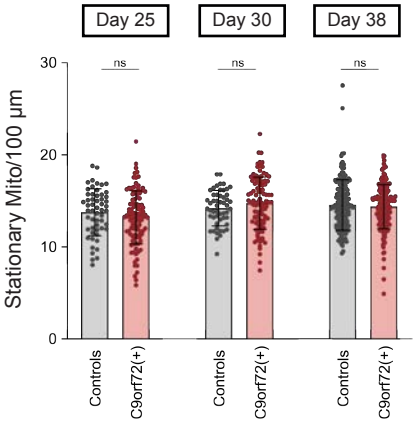

G

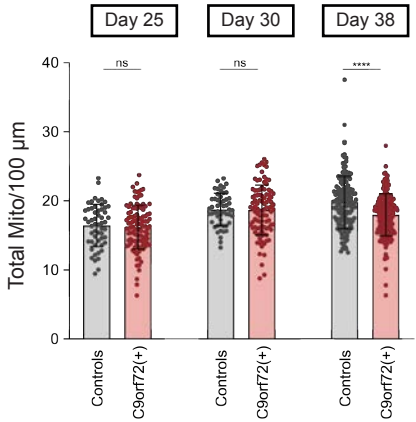

Fig.S7

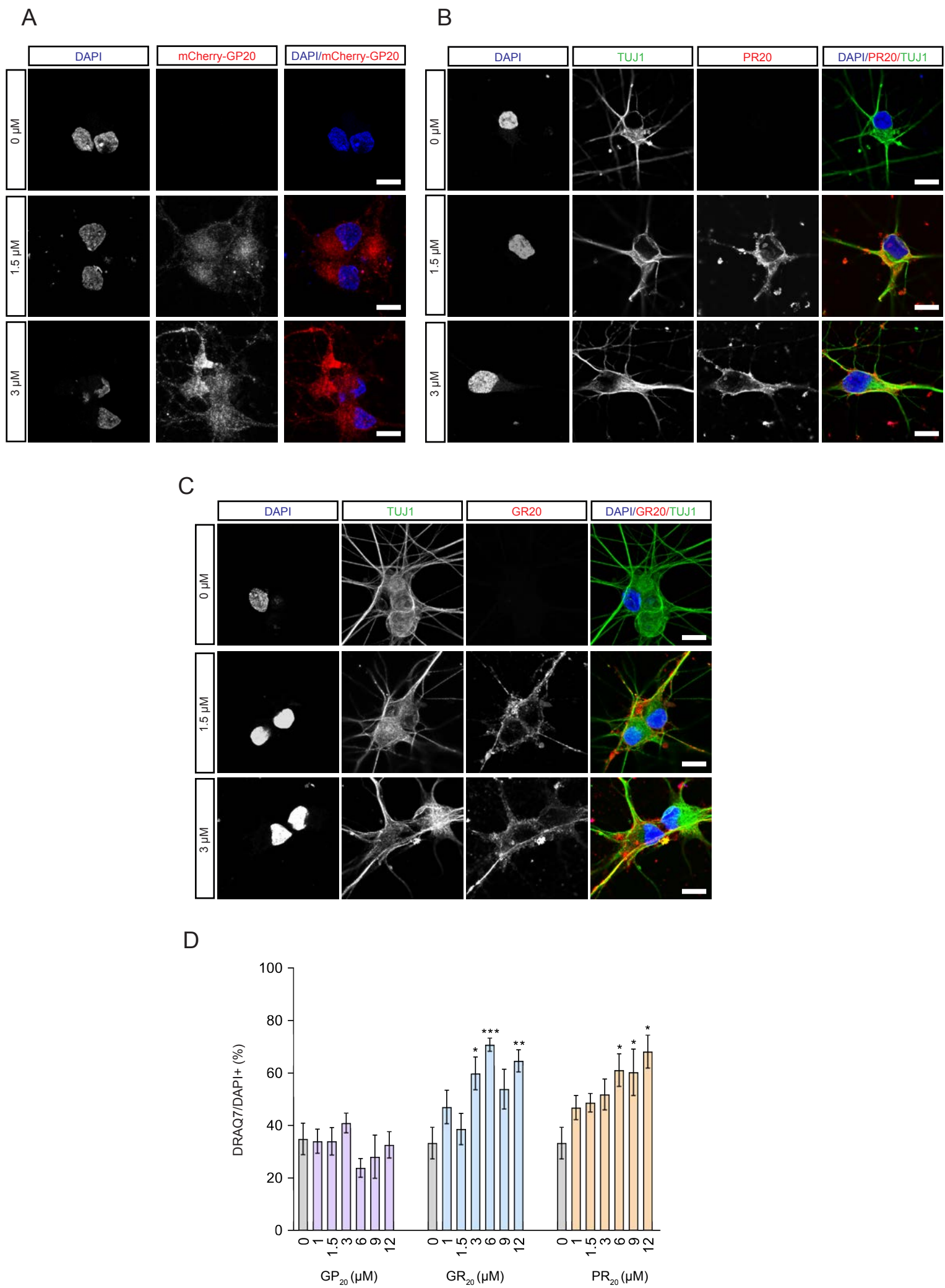

Fig.S8

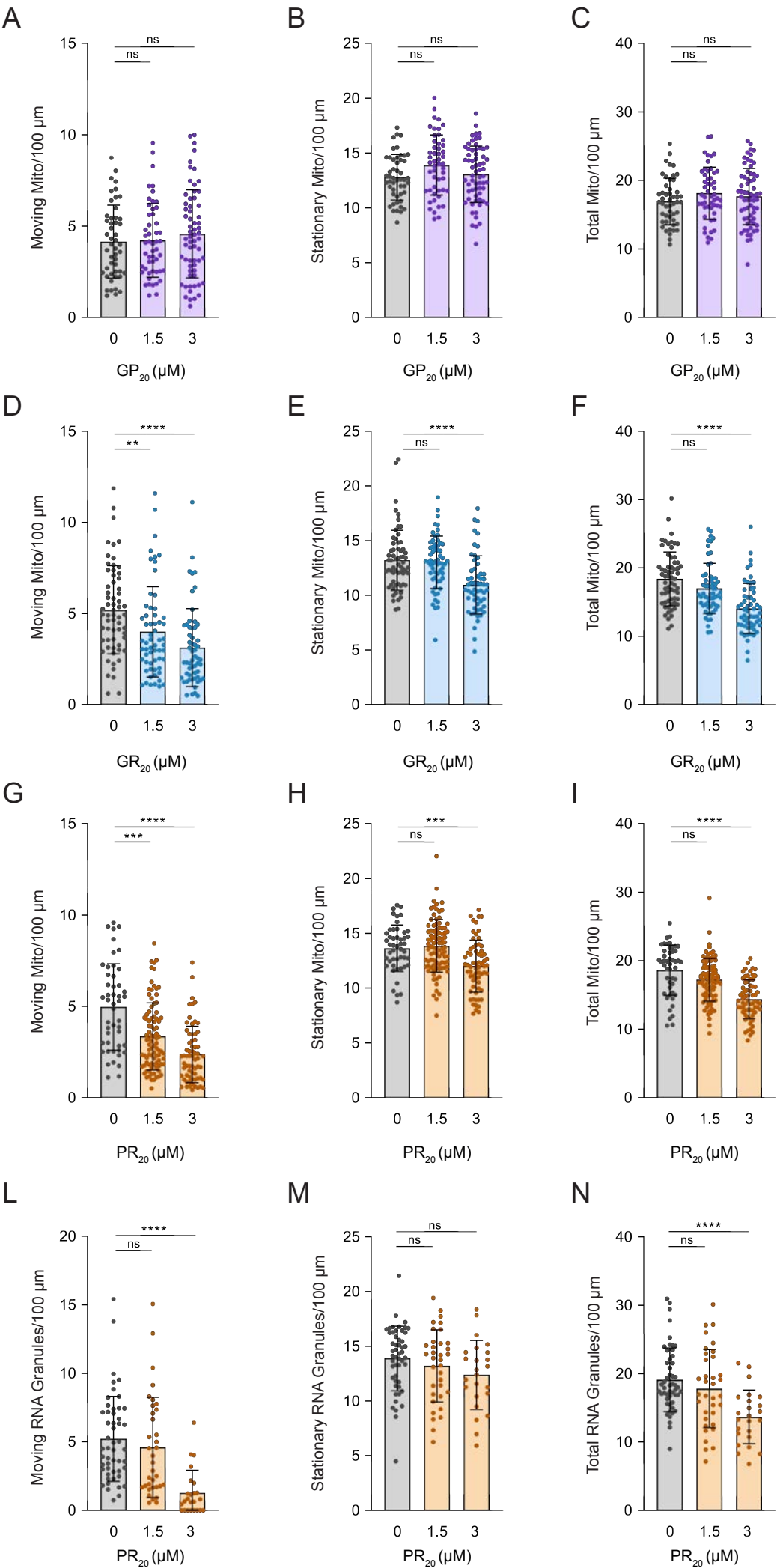

Fig.S9

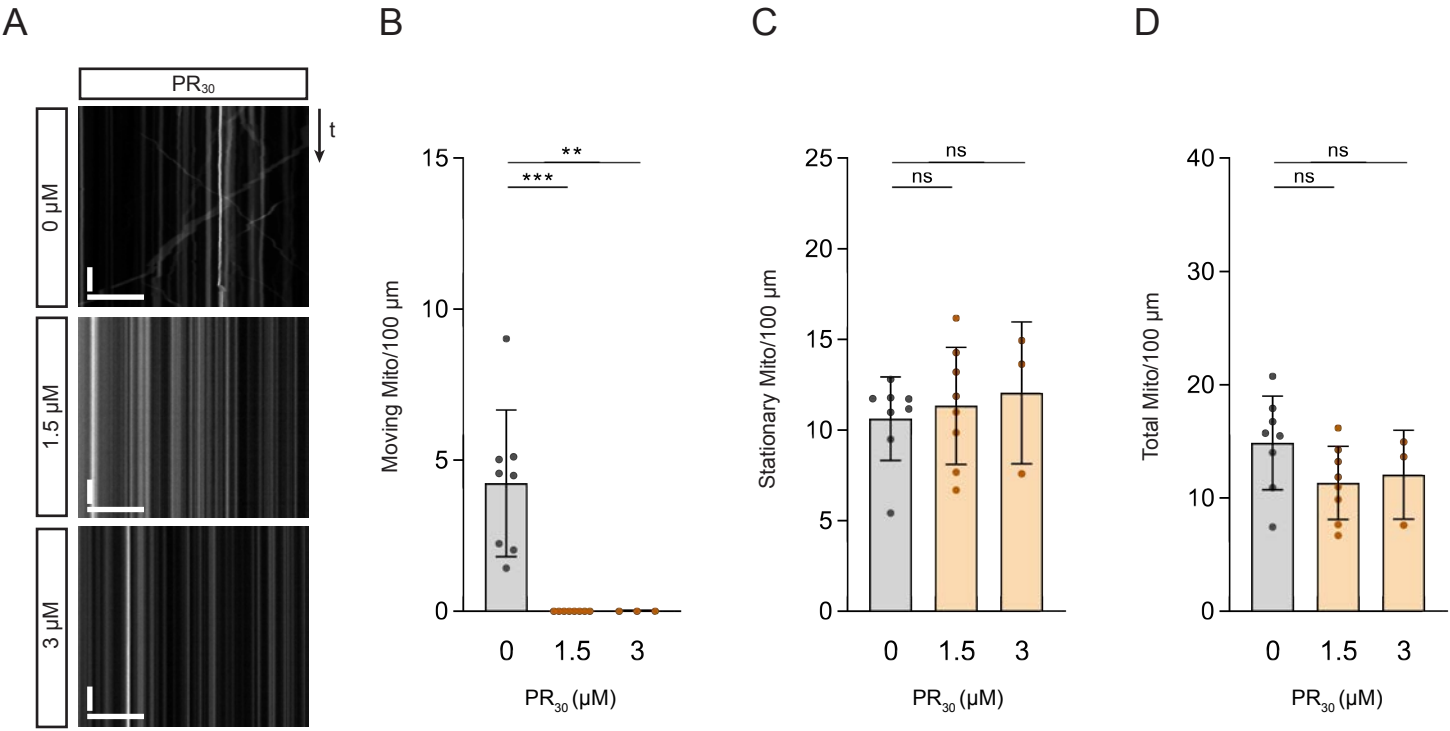

**Fig.S10**

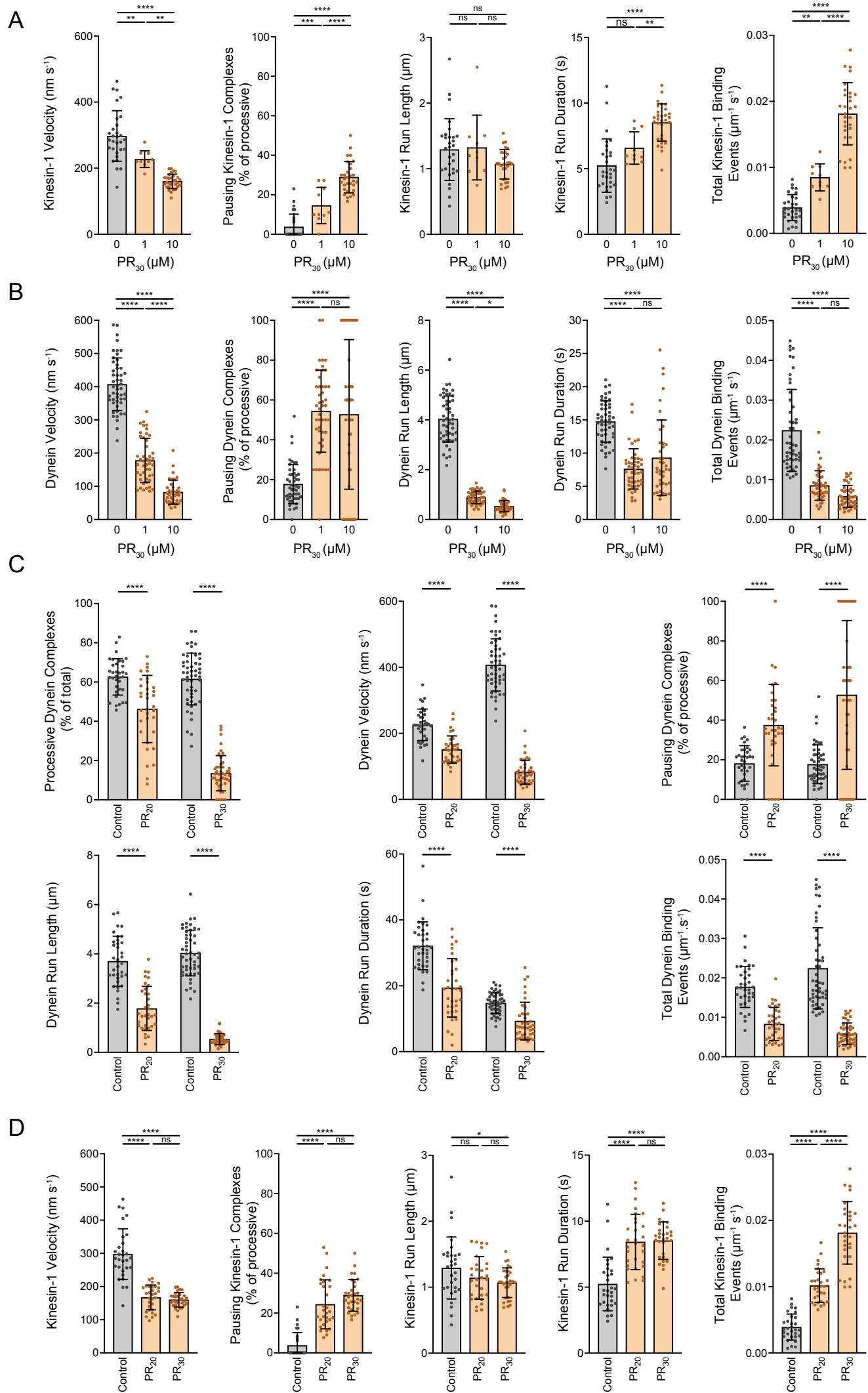

Fig.S11

A

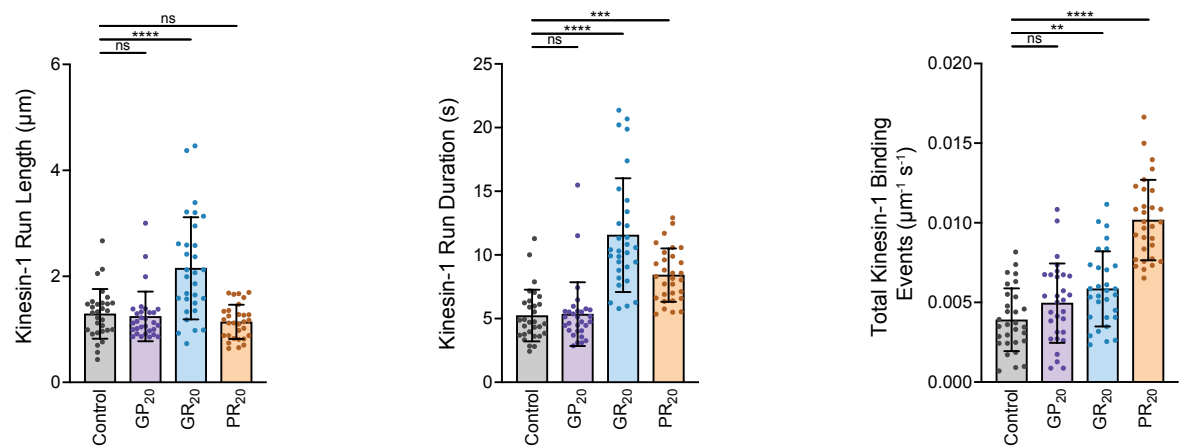

B

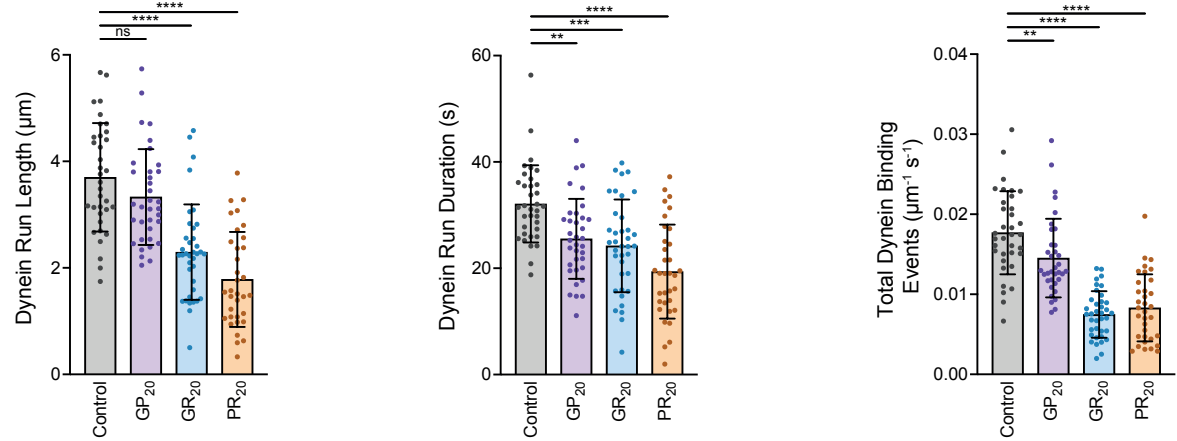

Fig.S12

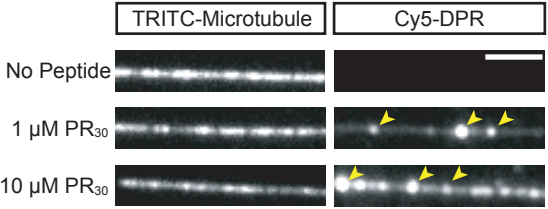

Fig.S13

A

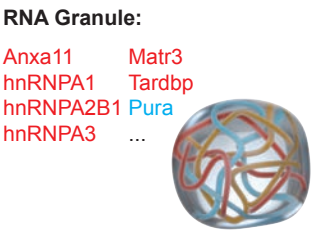

B

Other Cargoes:

|  |  |  |
| --- | --- | --- |
| GOCC | Mitochondrion | 1.29E-78 |
|  | RNP Complex | 1.68E-38 |
|  | Proteasome | 5.21E-26 |
|  | Synaptic Vesicle | 4.29E-18 |
|  | Endosome | 1.47E-09 |

C

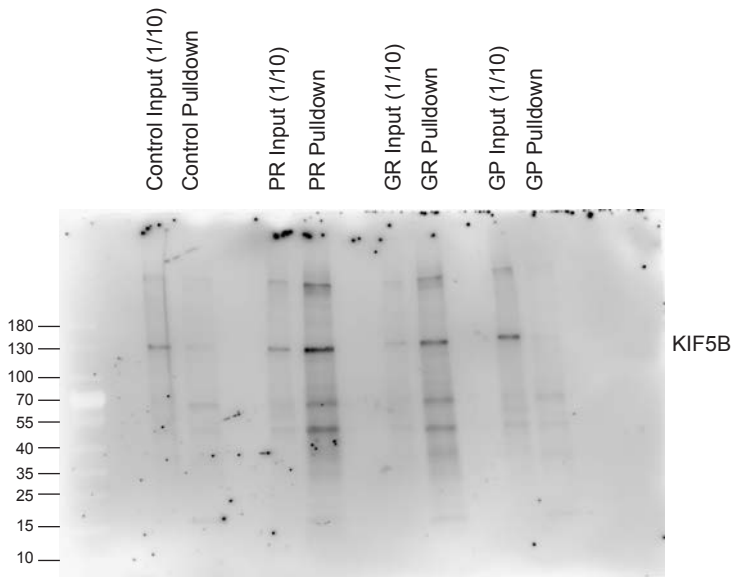

D

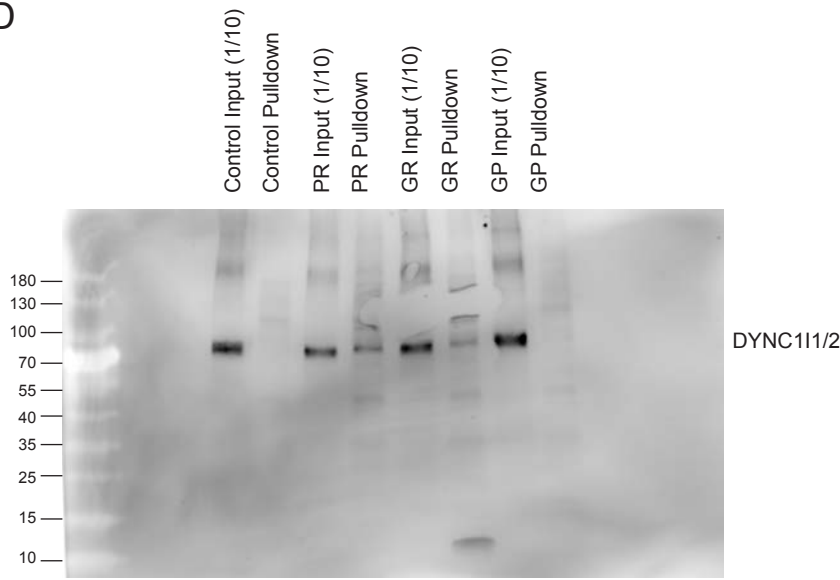

E

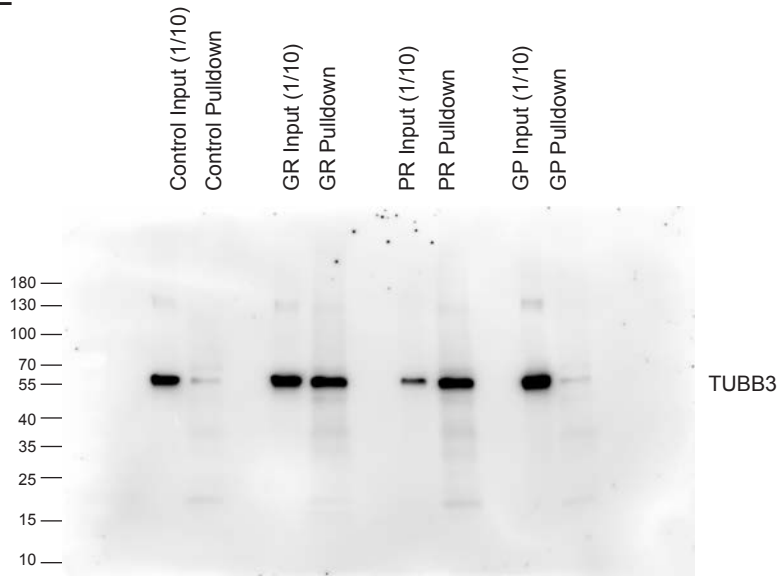

Fig.S14

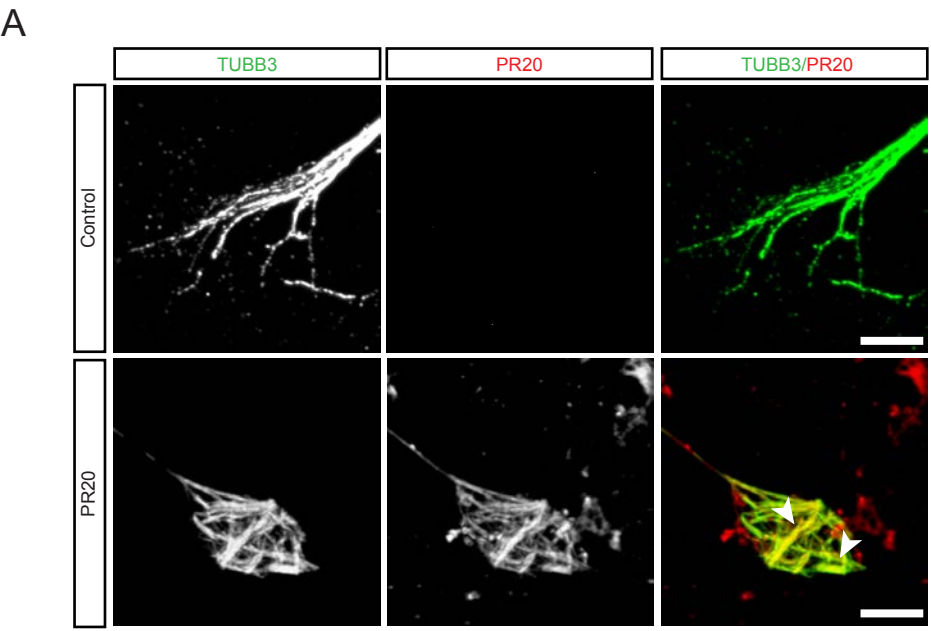

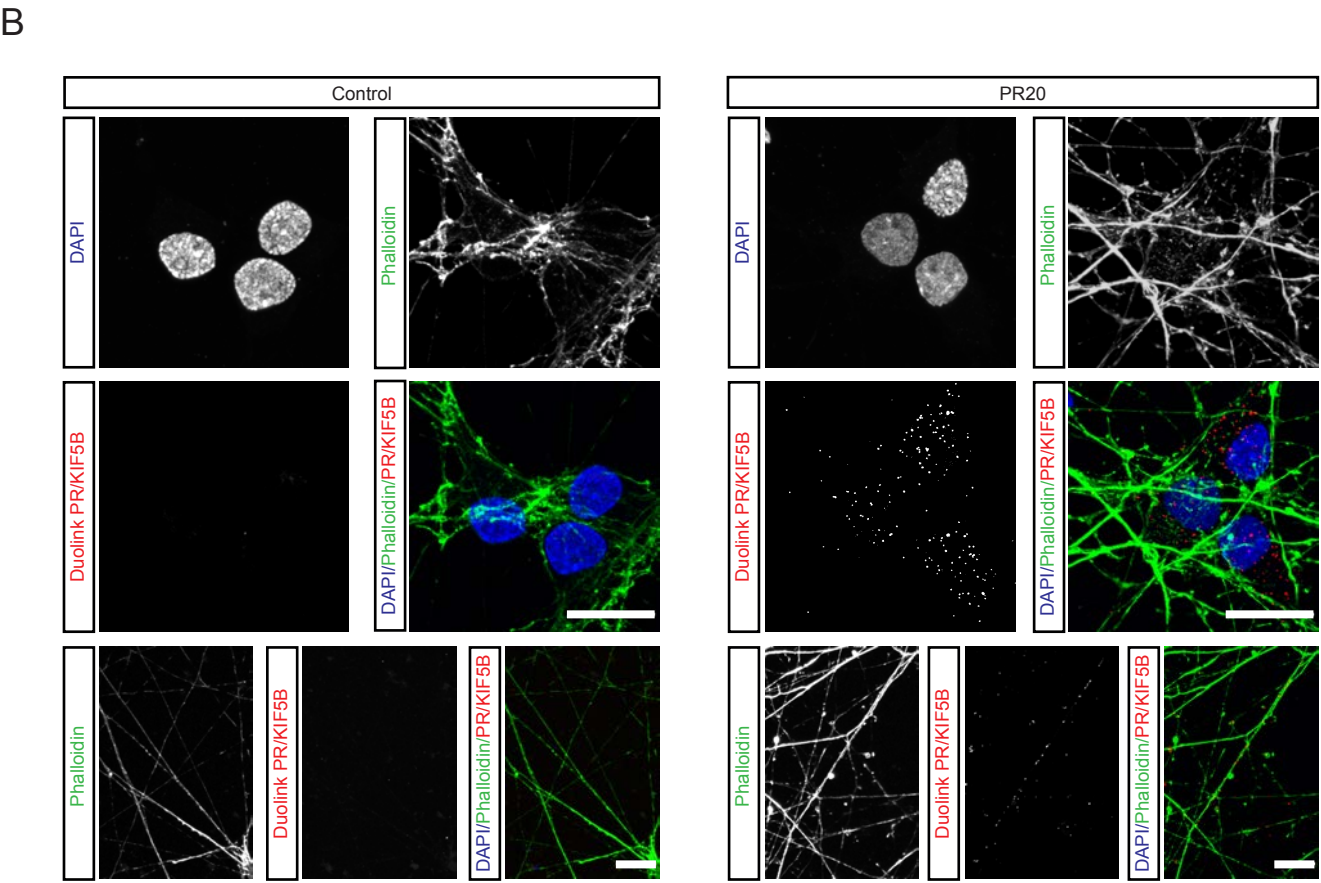

Fig.S15

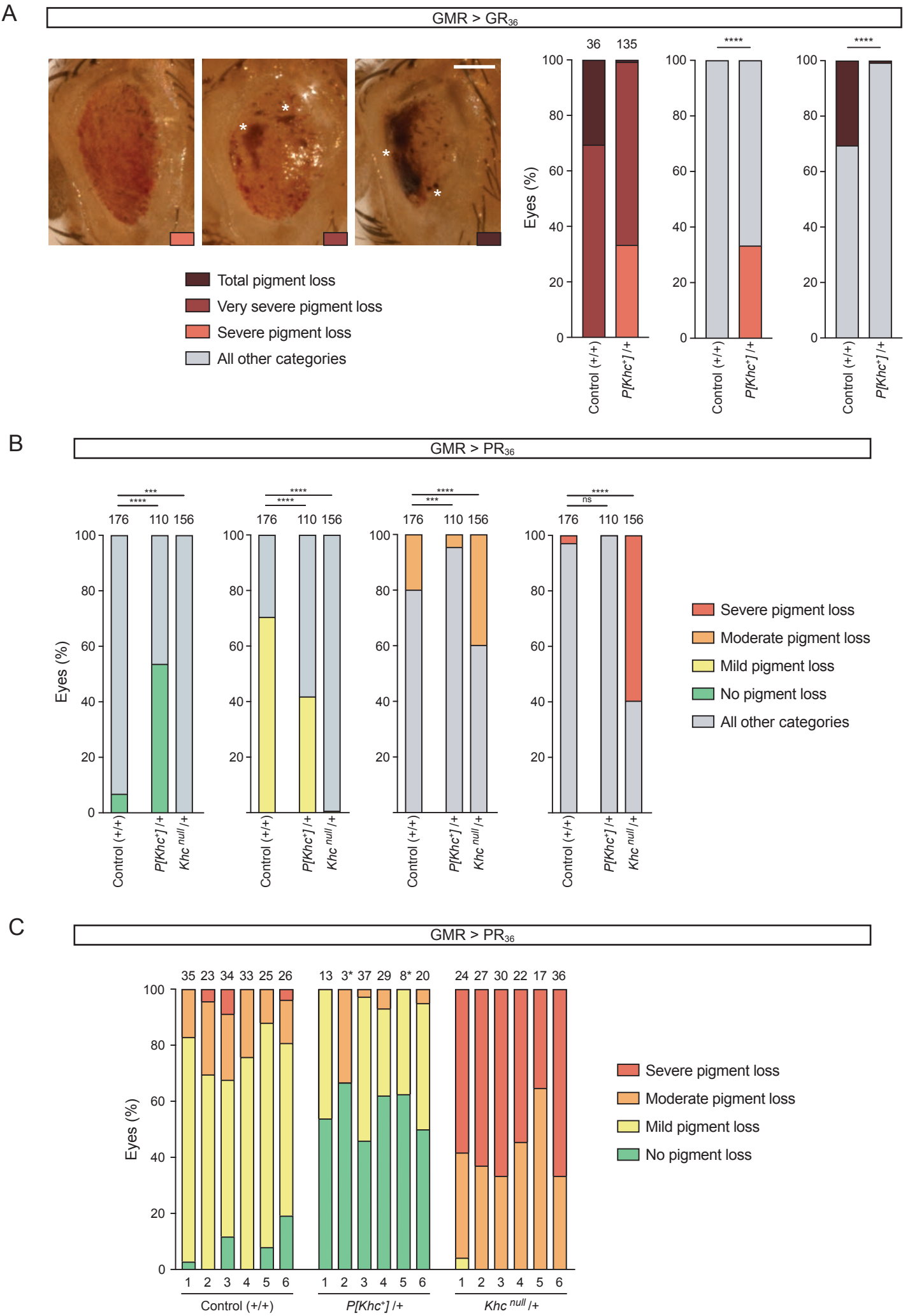
